# Loss of MGA mediated Polycomb repression promotes tumor progression and invasiveness

**DOI:** 10.1101/2020.10.16.334714

**Authors:** Haritha Mathsyaraja, Jonathen Catchpole, Emily Eastwood, Ekaterina Babaeva, Michael Geuenich, Pei Feng Cheng, Brian Freie, Jessica Ayers, Ming Yu, Nan Wu, Kumud R Poudel, Amanda Koehne, William Grady, A McGarry Houghton, Yuzuru Shiio, David P MacPherson, Robert N Eisenman

**Affiliations:** Basic Sciences Division, Fred Hutchinson Cancer Research Center, Seattle, WA; Human Biology and Public Health Sciences Divisions, Fred Hutchinson Cancer Research Center, Seattle, WA; Clinical Research Division, Fred Hutchinson Cancer Research Center, Seattle, WA; Comparative Pathology, Fred Hutchinson Cancer Research Center, Seattle, WA; Department of Medicine, University of Washington School of Medicine, Seattle, WA; Greehey Children’s Cancer Research Institute, The University of Texas Health Science Center, San Antonio, TX

## Abstract

MGA, a transcription factor and member of the MYC network, is mutated or deleted in a broad spectrum of malignancies. As a critical test of a tumor suppressive role, we inactivated Mga in two mouse models of non-small cell lung cancer using a CRISPR based approach. MGA loss significantly accelerated tumor growth in both models and led to de-repression of atypical Polycomb PRC1.6, E2F and MYC-MAX targets. Similarly, MGA depletion in human lung adenocarcinoma lines augmented invasive capabilities. We further show that MGA, E2F6 and L3MBTL2 co-occupy thousands of promoters and that MGA stabilizes PRC1.6 subunits. Lastly, we report that MGA loss has also a pro-growth effect in human colon organoids. Our studies establish MGA as a bona fide tumor suppressor in vivo and suggest a tumor suppressive mechanism in adenocarcinomas resulting from widespread transcriptional attenuation of MYC and E2F targets mediated by an atypical Polycomb complex containing MGA-MAX dimers.

## INTRODUCTION

Malignant progression often results from the de-regulation of factors that drive normal developmental programs. Given the critical role of transcription factors in driving cellular functions, it isn’t surprising that transcriptional regulators can function as potent tumor suppressors or oncogenes. Arguably, one of the most well-characterized oncogenic transcription factors is MYC, which is also essential for normal development (for reviews see Dang and Eisenman eds. 2014). Frequently altered across a broad spectrum of malignancies, MYC orchestrates transcriptional programs promoting metabolic re-wiring and ribosome biogenesis, ultimately facilitating rampant proliferation and tumor progression (reviewed in Stine et al. 2015). A large body of evidence suggests that MYC doesn’t function in isolation: it is part of a larger network of transcription factors that cooperate with or antagonize MYC activity (reviewed in Carroll et al. 2018). All members of this extended MYC network are basic-helix-loop-helix-leucine zipper (bHLHZ) transcription factors that recognize E-box motifs and other aspects of chromatin in thousands of genes. For example, MLX and MondoA can cooperate with MYC to promote tumorigenesis in MYC amplified cancers (Carroll et al. 2015). On the other hand, MXD 1-4 proteins, MNT and MGA, oppose MYC transcriptional activity by acting as repressors at E-boxes (Yang and Hurlin. 2017). In principle, either activation of MYC or inactivation of MXD function would stimulate expression of their gene targets and potentially promote oncogenesis. Some support for this notion is provided by recent TCGA data indicating that a subfraction of multiple tumor types are subject to deletions of MXD family proteins (Schaub et al. 2018). However, most striking are the genetic alterations sustained by the MAX dimerizing repressor, MGA, at high frequency across a wide range of cancers including 8% of lung adenocarcinomas (De Paoli et al. 2013, TCGA. 2014, Sun et al. 2017, Schaub et al. 2018). The majority of mutations in lung adenocarcinoma are truncating mutations (TCGA 2014) and enrichment in nonsense mutations at the MGA locus across all cancers is statistically significant, suggestive of a tumor suppressive role for the protein (Cooke et al. 2014). Consistent with this, MGA emerged as one of the top hits in a genome-wide CRISPR screen for tumor suppressors in DLBCL (diffuse large B-cell lymphoma) lines (Reddy et al. 2017).

We and others discovered and initially characterized MGA as a factor that specifically binds the small bHLHZ protein MAX (Hurlin et al.1999). MAX is the obligate dimerization partner of the MYC family of oncogenic drivers (Blackwood and Eisenman.1991, Amati et al. 1993,). MGA is unique in that it is a dual specificity transcription factor that contains both a TBX domain and a bHLHZ region. However, MGA is one of the least studied members of the network. Studies in mice and zebrafish suggest that MGA plays an important role in early development (Rikin and Evans. 2010, Sun et al. 2014, Washkowitz et al. 2015). Specifically, loss of MGA leads to peri-implantation lethality due to defects in polyamine biosynthesis (Washkowitz et al. 2015). Much of what is known of MGA’s function in mammalian cells results from studies in mouse embryonic stem cells (mESCs). An RNAi screen revealed that MGA and its dimerization partner MAX are both involved in repressing germ cell related transcripts (Maeda et al. 2013) and act to prevent entry into meiosis in germline stem cells by directly repressing meiotic genes (Suzuki et al. 2016). Moreover, a number of studies have shown that MGA interacts with atypical PRC1.6 complexes in ES cells and plays an integral role in recruiting PRC1.6 members to target promoters (Ogawa et al. 2002, Gao et al. 2012, Stielow et al. 2018, Scelfo et al. 2019, Fursova et al. 2019). Despite these advances in understanding MGA function in a developmental context, MGA’s role in tumorigenesis remains largely uncharacterized.

A recent study revealed that ectopic expression of MGA in lung adenocarcinoma cell lines retards their growth (Llabata et al. 2020). Here, we examine MGA’s function in tumors *in vivo* and dissect the molecular basis of its tumor suppressive activity. Given that MGA activity is lost at high frequency in lung adenocarcinoma, we utilized a lentiviral based *in vivo* CRISPR strategy to inactivate *Mga* in murine lung cancer models coupled with *in vitro* functional and genomic occupancy studies to elucidate the functional and molecular consequences. Importantly, we sought to characterize the role of MGA and PRC1.6 complexes in driving transcriptional repression in malignant settings. Lastly, we aimed to investigate a potentially broader function of MGA as a tumor suppressor by studying the functional consequences of MGA loss in normal and malignant colorectal cells.

## RESULTS

### *Mga* inactivation in *Kras*^G12D^ and *Kras*^G12D^*Trp53* −/− driven mouse models of lung cancer leads to accelerated tumorigenesis

We interrogated publicly available datasets in order to establish a correlation between MGA levels and tumor progression in lung cancer. We observed that MGA is mutated or deleted in 53 of 507 (10%) of lung adenocarcinoma patients (TCGA) and in 233 of 3696 (6%) of patients sequenced through the GENIE project (Fig. S1A). While MGA is a large gene, the majority of MGA mutations in both TCGA and GENIE datasets were truncating, suggestive of selection for inactivating mutations in tumorigenesis. In addition, low levels of MGA in lung adenocarcinoma significantly correlate with decreased overall survival (Fig. S1B, Győrffy et al. 2013). To ascertain whether *Mga* functions as a tumor suppressor *in vivo*, we utilized CRISPR-CAS technology to inactivate *Mga* in a *Kras*^LSL-G12D/+^ (henceforth termed Kras) driven mouse model of lung cancer (Jackson et al. 2001, Sánchez-Rivera. et al 2014). Intratracheal lentiviral delivery of Cre, Cas9 and sgRNA against *Mga* using the lentiCRISPRv2 cre vector (Walter et al. 2017) (Fig. 1A) results in the activation of *Kras* ^G12D^ along with indel formation at the *Mga* locus (Fig. S1C). For survival analysis, mice were euthanized when moribund from lung tumor burden. There was a significant decrease in survival in *Kras* ^G12D^ mice receiving sg*Mga* when compared to empty vector treated controls (Fig. 1B). Tumors from sg*Mga* treated mice exhibited a higher proliferation index as ascertained by Ki67 staining when compared to empty controls (Fig. 1C, 1D). Pathological examination of lesions at endpoint by a board certified veterinary pathologist (A.K.) revealed that mice in both groups had a spectrum of grade 1-5 tumors characteristic of lesions that develop in the Kras model (Fig. S1D, S1E) (Jackson et al. 2005). Sequencing of tumors confirmed the presence of indels predicted to result in premature truncation in 75% (6/8) tumors isolated from sgMga treated mice. We next examined the potential synergy of *Mga* loss with the inactivation of other tumor suppressors. Intriguingly, TCGA and GENIE data revealed that there is a significant co-occurrence of genomic alterations in *MGA* and *TP53* in lung adenocarcinoma patients (Fig. S1F). To further investigate putative cooperation between *MGA* and *TP53 loss*, we utilized the *Kras*^LSL-G12D/+^*Trp53*^fl/fl^ (henceforth termed KP) mouse model of lung cancer. We inactivated *Mga* in this model using the same lentiCRISPRv2 cre approach described previously and analyzed mice at a common time point 3 months later. At 3 months post intra-tracheal instillation of lentiCRISPRv2cre, KP-sg*Mga* mice had a substantially higher tumor burden when compared to KP-Empty controls (Fig. 1E, 1F). This was accompanied by an overall increase in number of tumors and tumor grade upon Mga inactivation (Fig. 1G, 1H). Immunostaining revealed that sg*Mga* tumors had an increased proportion of Ki67+ tumor cells when compared to controls (Fig. 1I, 1J). Consistent with increase in tumor grade, sg*Mga* tumors had increased vasculature (as evidenced by MECA32 staining) when compared to controls (Fig. S1G, S1H). In addition to analyzing mice at a common 3-month time point, we also aged a separate small cohort of infected animals to isolate tumors and generate cell lines. These cell lines (herein designated KP cells) were utilized to further characterize sg*Mga* inactivated tumors. A substantial decrease in MGA protein levels was observed in most KP-sgMga cell lines (Fig. 1K,1L, S1I). We confirmed that KP-sg*Mga* lines are dependent on the inactivation of *Mga* as ectopic expression of MGA in KP-sg*Mga* lines impaired proliferation (Fig 1M,1N).

**Figure 1:**
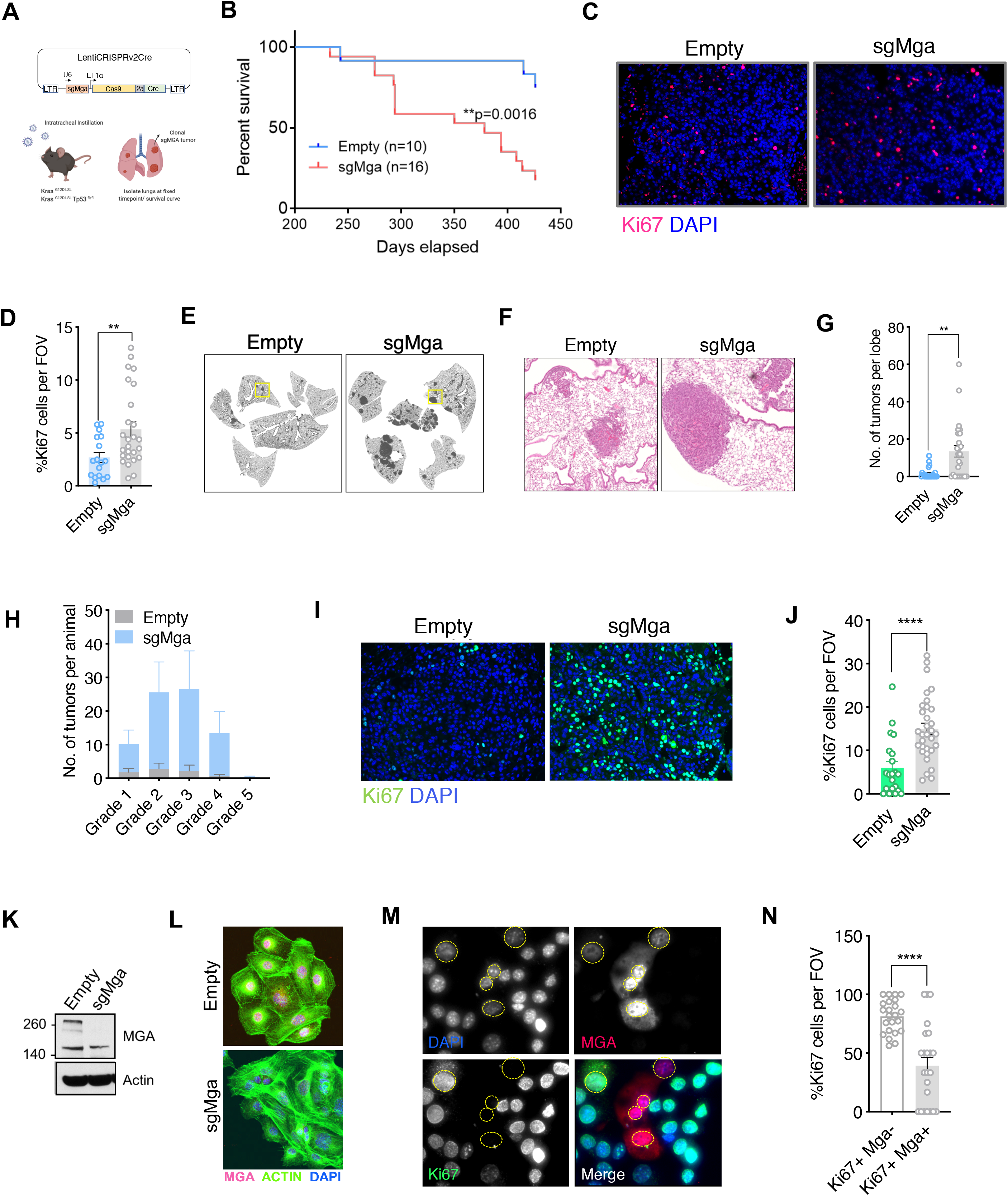
*Mga* inactivation *in vivo* leads to accelerated tumorigenesis. (A) Schematic of lentiCRISPRv2cre vector and strategy used for in vivo Kras and KP experiments. (B) Kaplan-Meier curve for survival in sgMga vs Empty virus treated Kras mice (p-value computed using Log rank test). (C) Representative micrographs and, (D) quantification of Ki67 immunostaining on Kras lung tumor tissue from sgMga and Empty control mice (n=5 mice per group. **p<0.01. Calculated using Welch’s t-test assuming unequal variance). (E) Representative H&E staining of sgMga and Empty lungs from KP mice harvested 3 months post-intratracheal instillation. (F) Higher magnification of selected regions from (E) showing individual tumors in both groups. Quantification of (G) number of lesions per lobe and, (H) number of lesions per grade in Empty and sgMga animals (n=5 mice per group. **p<0.01). (I) Representative images and, (J) quantification of Ki67 staining to assess proliferation in KP Empty and KP sgMga lung tumors harvested 3 months post-infection (n=32 tumors from 4 sgMga mice n=21 tumors from 2 Empty mice. ****p<0.0001. Calculated using Welch’s t-test assuming unequal variance). (K) Western blot and, (L) immunofluorescent staining for MGA in cell lines derived from KP tumors. (M) Representative images and (N) quantification of Ki67 and MGA co-staining in an sgMga KP line upon ectopic expression of MGA (n= 23 FOV each from 3 independent experiments. ****p<0.0001). All p-values calculated using a 2-sided Student’s t-test unless otherwise noted. Error bars represent SEM.

### MGA loss de-represses PRC1.6 and MYC targets and up-regulates pro-invasion genes

To identify transcriptional changes that may underly the phenotypes we observed, we performed RNA-seq analyses. We compared gene expression profiles between 6 Kras sg*Mga* and 8 control Kras lung tumors (Fig. S2A). sg*Mga* tumors were sequenced and only those confirmed to have indels were used for RNA-seq experiments. Principal component analysis suggested heterogeneity in gene expression across tumors (Fig. S2B). This may be due to varying levels of non-tumor cells such as immune cells and fibroblasts present within tumors in this model (Busch et al. 2016). Despite the heterogeneity, Mga inactivated cells exhibited a striking increase in expression of several meiotic genes, previously reported to be targeted by MGA-PRC1.6 in ES and germline cells, such as *Stag3, Sox30* and *Tdrd1* (Fig. 2A) (Suzuki et al. 2016). This suggests that similar to ES cells, MGA also assembles into PRC1.6 complexes in somatic normal and tumor cells. In agreement with our Ki67 data, *Ccnd1* levels were elevated in *sgMga* tumors compared to control (Fig. 2A). Gene set enrichment analysis revealed that several metabolic pathways and a subset of MYC-regulated genes were also upregulated in sg*Mga* tumors, indicating that MGA represses MYC targets, similar to repression by MXDs seen in other tumor models (Fig. 2B) (Yang and Hurlin. 2017). Intriguingly, inactivation of *Mga* led to the downregulation of genes involved in anti-tumor responses, such as NK cell markers and interferon signaling genes (Fig. 2C; Fig. S2C). This is also consistent with a role for MYC in repressing anti-tumor immune responses in tumors, including in Kras driven lung and pancreatic adenocarcinoma (Casey et al. 2016, Kortlever et al. 2017, Muthalagu et al. 2020).

**Figure 2:**
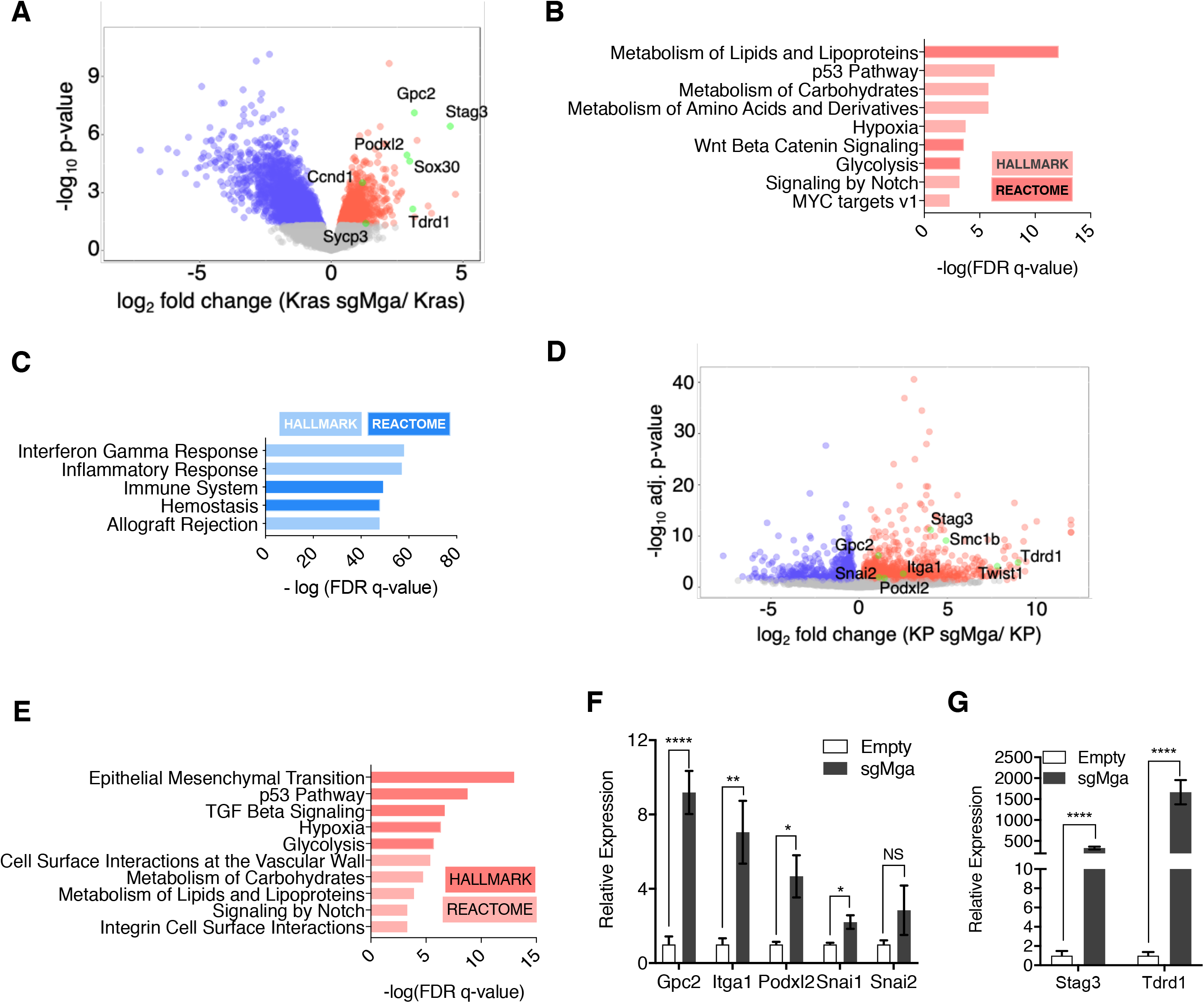
MGA loss leads to the de-repression of PRC1.6 and MYC targets and up-regulation of pro-invasion genes. (A) Volcano plot of Kras tumor RNA profiling data comparing Kras and Kras sgMga tumors. Upregulated pro-invasion and PRC1.6 targets highlighted in green. Hallmark and Reactome GSEA analysis for pathways enriched for in (B) upregulated and, (C) downregulated genes in Kras sgMga tumors compared to empty controls. (D) Volcano plot of KP cell line RNA Seq data showing PRC1.6 and pro-invasive genes in green (n= 2 cell lines in triplicate for KP control and 3 cell lines in triplicate for KP sgMGA). (E) Hallmark and Reactome analysis of genes upregulated in KP sgMga vs. Empty cell lines. qPCR on Empty and sgMga KP lines to confirm levels of (F) pro-invasive and, (G) PRC1.6 meiotic targets (n=6 for each – 2 sets of RNA from 3 independent lines for sgMga and Empty). *p<0.05, **p<0.01, ****p<0.0001. All p-values calculated using a 2-sided Student’s t-test. Error bars represent SEM.

Next, we performed RNA-Seq on MGA inactivated and control KP lines. Cell lines were used to overcome the limitation of having a mixed population of cells in primary tumors as with the Kras tumors. Despite the differences in genetic background (Kras vs. KP), several PRC1.6 complex repressed germ cell related transcripts, including *Stag3*, were highly upregulated upon MGA loss in KP cells, similar to Kras tumors (Fig 2D). In addition, we also observed an upregulation of pro-invasive genes such as *Podxl2, Gpc2, Snai2* and *Itga1* (Fig. 2D). *Gpc2 and Podxl2* also appeared to be increased in Kras sgMga tumors (Fig. 2A). Gene set enrichment analysis revealed that Epithelial to Mesenchymal transition (EMT) and TGF-beta signaling were amongst the most significantly enriched pathways in sg*Mga* KP cells (Fig 2E). Intriguingly, genes upregulated in sgMga KP cells correlated significantly with those up-regulated upon MYC, E2F4 and TWIST1 overexpression in other systems (Fig. S2D). In addition, there was also a significant correlation with genes reported to be up-regulated upon PCGF6 knockdown or MYC overexpression (Fig. S2D). Along with the increase in tumor grade and MECA32 staining we observed in the KP model, these results corroborate a role for MGA in repressing invasion (Fig. S1D, S1G and see below). We noticed some commonalties in pathways de-repressed upon MGA loss in Kras and KP tumors. Processes such as hypoxia, glycolysis, carbohydrate and lipid metabolism found to be enriched for in sg*Mga* Kras tumors also scored significantly in the KP cells (Fig. 2B, 2E). However, as determined by gene set enrichment, there wasn’t a major overlap for genes downregulated in sgMga cells (Fig. S2E). We further confirmed the activation of EMT and pro-invasive genes like *Snai1, Itga1, Podxl2 and Gpc2* (Fig. 2F) and PRC 1.6 targets *Stag3* and *Tdrd1* (Fig. 2G) via qPCR on a panel of control and sgMga KP lines.

Taken together, our data suggest that MGA regulates meiotic transcripts such as *Stag3* and MYC targets in both ES cells and lung tumor initiating cells but also regulates distinct gene signatures based on cellular context, such as *Trp53* expression.

### MGA loss correlates with activation of *STAG3* AND *PODXL2* in human lung cancer and results in a pro-invasive phenotype *in vitro*

We next asked if the phenotypic and expression changes we observed in our *in vivo* studies are also manifested in patients and established human lung cancer cell lines. Given the overlap of MGA targets we observed in our mouse models, we examined human lung adenocarcinoma TCGA data to assess whether the expression of these genes correlates with *MGA* loss in patients. We then grouped patients into MGA WT and MGA altered groups based on the presence of mutations and deletions at the MGA locus to assess functional enrichment of expression of genes in MGA altered tumors. Interestingly, STAG3 appeared to be highly expressed (SD > 3 above mean) in a significant subset of MGA altered cases (truncating mutations and deletions) (Fig. S3A, S3B). Strikingly, genes overexpressed in the MGA altered group are significantly enriched for MYC, MAX, E2F6 and E2F4 in Enrichr ChEA analysis (Fig. S3C). Consistent with mouse Kras tumor data, there appeared to be a significant downregulation of anti-tumor immune responses and decrease in IFN signaling genes such as *IFITM1* in MGA altered cases (Fig. 3A, 3B, S3D). Importantly, functional enrichment analysis on cases with MYC amplifications vs. non-amplified revealed a similar trend with regards to anti-tumor immune responses and IFN signaling (Fig. S3E). We then sought to overlap genes upregulated in our mouse Kras and KP expression profiling data (>1.5X fold change, adj. p-value <0.05) with genes overexpressed in the MGA altered TCGA group (p-value <0.05). Amongst the 5 gene overlap, we observed that established PRC1.6 target STAG3 and pro-invasion gene PODXL2 were significantly altered in all 3 comparisons (Fig. 3C,3D,3E).

**Figure 3:**
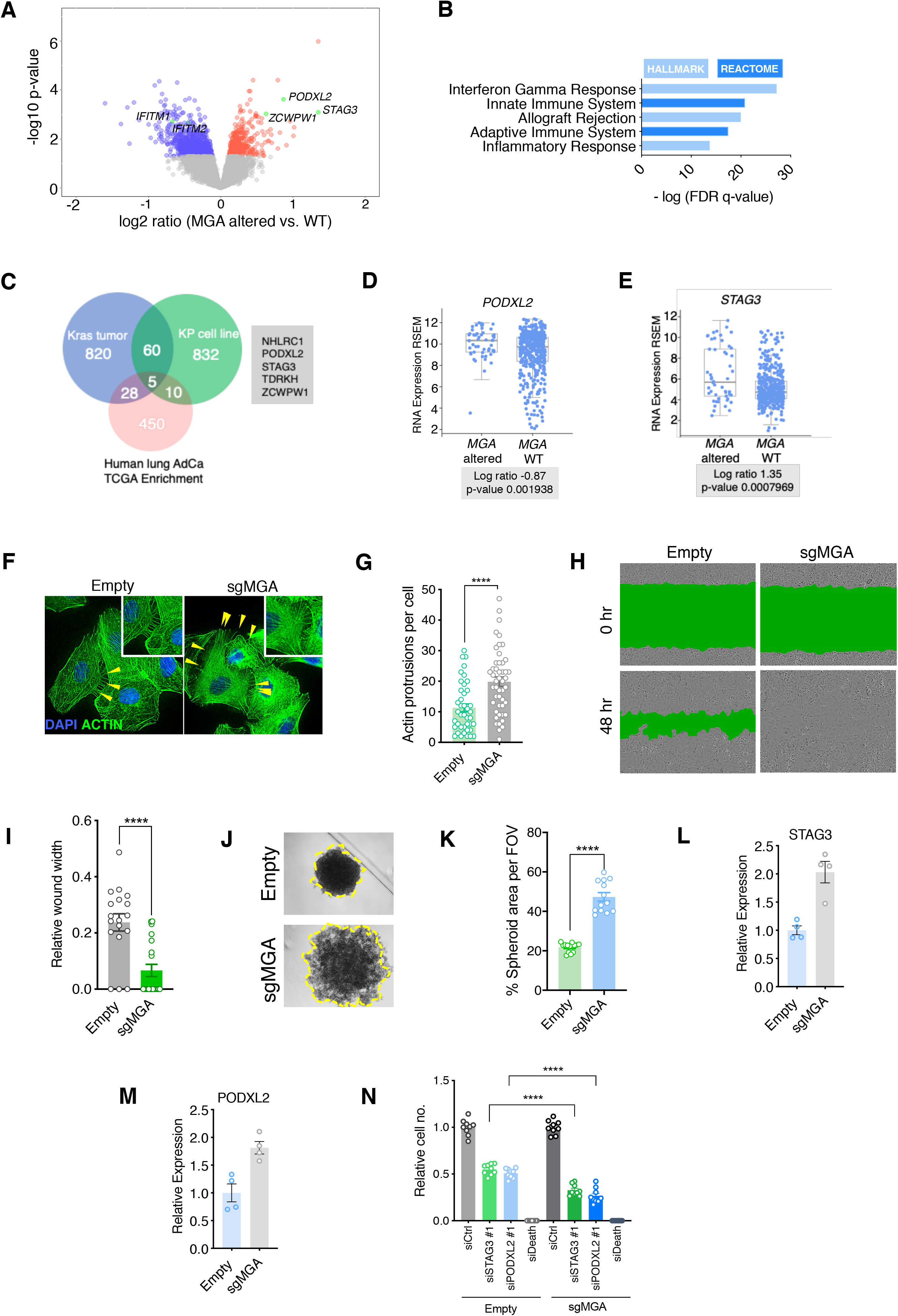
MGA loss correlates with activation of *STAG3* AND *PODXL2* in human lung cancer and results in a pro-invasive phenotype *in vitro.* (A) Volcano plot of transcripts that are differentially expressed in MGA altered (n=52) vs. WT (n=455) pan TCGA lung adenocarcinoma data. Representative genes highlighted in green. (B) Hallmark and Reactome analysis of genes downregulated in MGA altered patients vs. non-altered (p-value <0.05). (C) Venn diagram depicting overlap between genes upregulated in the mouse Kras tumor, KP cell line and Human lung TCGA data with MGA alterations. Log ratio of (D) PODXL2 and, (E) STAG3 expression in MGA altered vs. WT lung adenocarcinoma patients (TCGA, 2014). (F) Representative Phalloidin (Actin) staining and (G) Quantification of actin protrusions in Empty and sgMGA A549 cells (n=41 cells Empty and n=48 cells sgMGA. 4 independent experiments for each group. ****P<0.0001). (H) Representative wound widths and, (I) quantification of wound width at 48 hours in Empty and sgMGA wells (n= 18 Empty n=20 sgMGA. ****P<0.0001). (J) Spheroid formation and (K) quantification in Empty and sgMGA A549 lines (quantification at Day 6, n=12 spheroids for each from 3 independent experiments. ****P<0.0001). qPCR for (L) PODXL2 and (M) STAG3 in Empty and sgMGA A549 cells (n=4 for each condition). (N) Cell growth upon siRNA knockdown of STAG3 and PODXL2 in A549 Empty and sgMGA cells. All p-values calculated using a 2-sided Student’s t-test unless otherwise noted. Error bars represent SEM.

To investigate whether *MGA* loss leads to similar functional consequences *in vitro*, we used either shRNA or CRISPR based approaches to delete or knock down *MGA* in A549 and NCI-H23 lung adenocarcinoma lines (Fig S3F, S3H). Interestingly, MGA suppression conferred no growth advantage to these lines, in contrast to the phenotypes we observed in models where MGA was inactivated at the tumor initiation stage (Fig S3G, S3I; compare with Fig. 1B, 1F, 1I). However, we observed an increase in invasive properties upon MGA inactivation, as evidenced by a significant increase in the number of actin protrusions present on cells (Fig. 3F, 3G; Fig. S3J,K). This was accompanied by faster migration of MGA inactivated cells in wound healing assays (Fig. 3H,3I, S3L, S3M) and formation of less compact spheroids by A549 sgMGA cells (Fig 3J, 3K). In addition, we observed an up-regulation of *PODXL2* and *STAG3* (Fig. 3L,3M). In order to study the importance of these genes in lung cancer cells, we performed siRNA knockdowns of STAG3 and PODXL2 in control and sgMGA cells (Fig S3N, S3O). Knockdown of either gene led to a substantial reduction in cell growth and this effect was more pronounced in sgMGA cells (Fig. 3N). Taken together, these results indicate that the changes we observed in mouse models, including the up-regulation of MYC and PRC1.6 targets upon MGA loss, are consistent with those observed in patient data and human lung cancer lines.

### MGA stabilizes PRC1.6 complex members in lung cancer cells

Given the consistent up-regulation of *STAG3* and *PODXL2* in Kras tumors, KP tumor cells and human lung cancer lines upon MGA inactivation (Fig. 2A, 2F, 2G Fig. 3 A, 3C-E), we sought to identify proteins that interact with MGA and mediate its function as a repressor. We performed tandem affinity purification of double tagged (FLAG and HIS) MGA in 293T cells followed by mass spectrometry to identify MGA interactors. We confirmed that full length MGA or MGA fragments (aa967-1300 and aa2153-2856) interact with several repressive molecules and complexes (protein prophet score > 0.9). These include members of the PRC1.6 complex such as RING2 and L3MBTL2 (Fig. 4A). In addition, MGA was also seen to bind members of the chromatin modifier MLL-WRAD complex and HDACs 1 and 2 (Fig. 4A). Studies in ES cells suggest that MGA is required for PRC1.6 assembly and stabilizes some components of the complex (Gao et al. 2012, Stielow et al. 2018, Scelfo et al. 2019). Therefore, we determined levels of PRC1.6 members MAX, L3MBTL2 and E2F6 in *Mga* inactivated and control KP lines. Mga loss resulted in decreased protein levels of E2F6, PCGF6 and L3MBTL2 (Fig. 4B-E) despite RNA levels remaining fairly constant (Fig. S4) whereas MAX levels remain unchanged (Fig. 4B, Fig. S4). We further confirmed a decreased E2F6, L3MBTL2 and PCGF6 protein levels in *MGA* mutant human non-small cell lung cancer lines H2291 and Lou-NH-91 when compared to wildtype lines such as NCI-H1975, NCI-H23 and 91T (Fig. 4F-I). Overall, our data suggest that MGA functions in maintaining the integrity of the PRC1.6 complex even in malignant cells.

**Figure 4:**
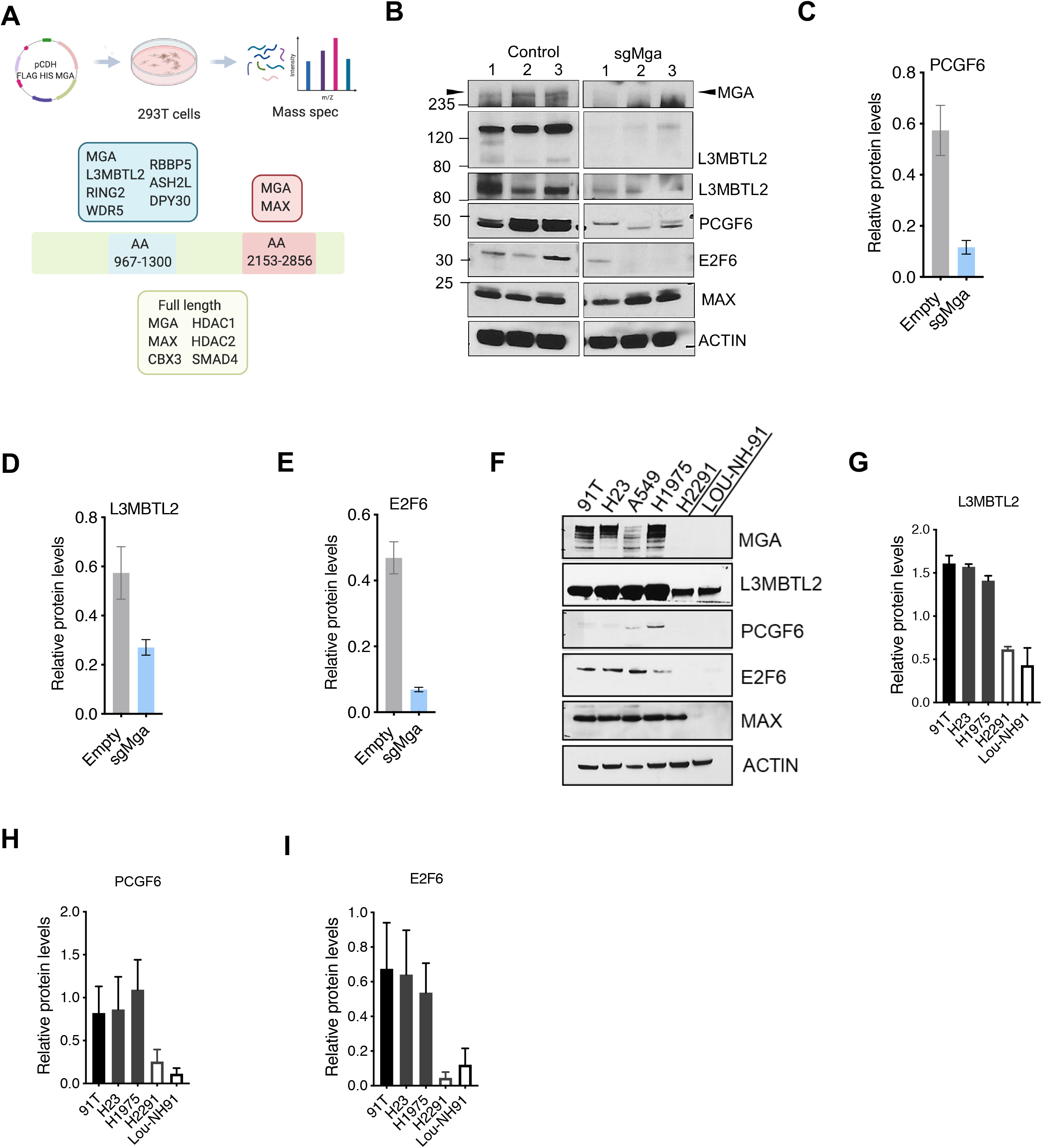
MGA stabilizes PRC1.6 complex members in lung cancer cells. (A) Mass spectrometric analysis of interactors of full length, aa967-1300 and aa2153-2853 fragments of FLAG-His-double tagged MGA. Confirmed interactors shaded in color (protein prophet score >0.9). (B) Representative immunoblot to show protein levels of MGA, L3MBTL2 (lower panel to show L3MBTL2 at higher exposure), PCGF6, E2F6 and MAX in control and sgMga mouse KP lines (n=3 individual cell lines for control and sgMga). Densitometry of PRC1.6 members (C) L3MBTL2 (D) PCGF6 and (E) E2F6 in control and sgMGA KP lines. (F) Representative immunoblots for MGA, L3MBTL2, PCGF6, E2F6 and MAX in MGA WT and mutant human non small cell lung cancer lines. Quantification of protein levels of (G) L3MBTL2 (H) PCGF6 and (I) E2F6 in MGA WT (blacks bars) vs. MGA mutant (white bars) lung cancer lines. For densitometry, 3 independent western runs were used for each comparison. Error bars represent SEM.

### MGA is a determinant of PRC1.6 genomic binding in tumor cells

Genomic targets of the PRC1.6 complex have been identified in select cellular contexts, including in 293FT cells and mouse ES cells (Gao et al. 2012, Stielow et al. 2018, Scelfo et al. 2019). To assess the direct role of PRC1.6 complexes in mediating repression in tumor cells, we performed conventional ChIP-Seq (X-ChIP: crosslinked chromatin immunoprecipitation followed by sequencing). We interrogated MAX, MGA, L3MBTL2 and E2F6 occupancy in a *Mga-* inactivated and control KP line. MGA, MAX, E2F6 and L3MBTL2 were seen to bind thousands of promoters in MGA expressing KP tumor cells suggesting that the PRC1.6 complex may play a broad role in tumor cell gene expression (Fig. 5A). Strikingly, we observed a substantial overlap between MGA, MAX, E2F6 and L3MBTL2 binding in control KP cells (Fig. 5A). Moreover, we noted a marked reduction in L3MBTL2 and E2F6 binding in KP-sgMga cells whereas MAX binding was largely unaffected (Fig. 5B, 5C, Fig. S5A). These binding data could reflect MGA’s role in both recruitment and stabilization of PRC1.6 complex members. Specifically, we saw significant enrichment of MGA binding at several genes that are up-regulated in sgMga KP cells, such as *Stag3* and *Podxl2* (Fig. 5D, Fig. 5F). Enrichr analysis of MGA bound genes revealed a significant enrichment for MYC, MAX and E2F (E2F4) bound genes (Fig. 5E). Of note, MYC and MAX binding at promoters in MGA null cells is largely unperturbed and even increased in certain cases (Fig. S5A). For example, MYC binding is augmented at *Stag3* and *Podxl2* in sgMga cells, in contrast to decreased occupancy of L3MBTL2 and E2F6 at the same loci (Fig. 5F, Fig. S5B).

**Figure 5:**
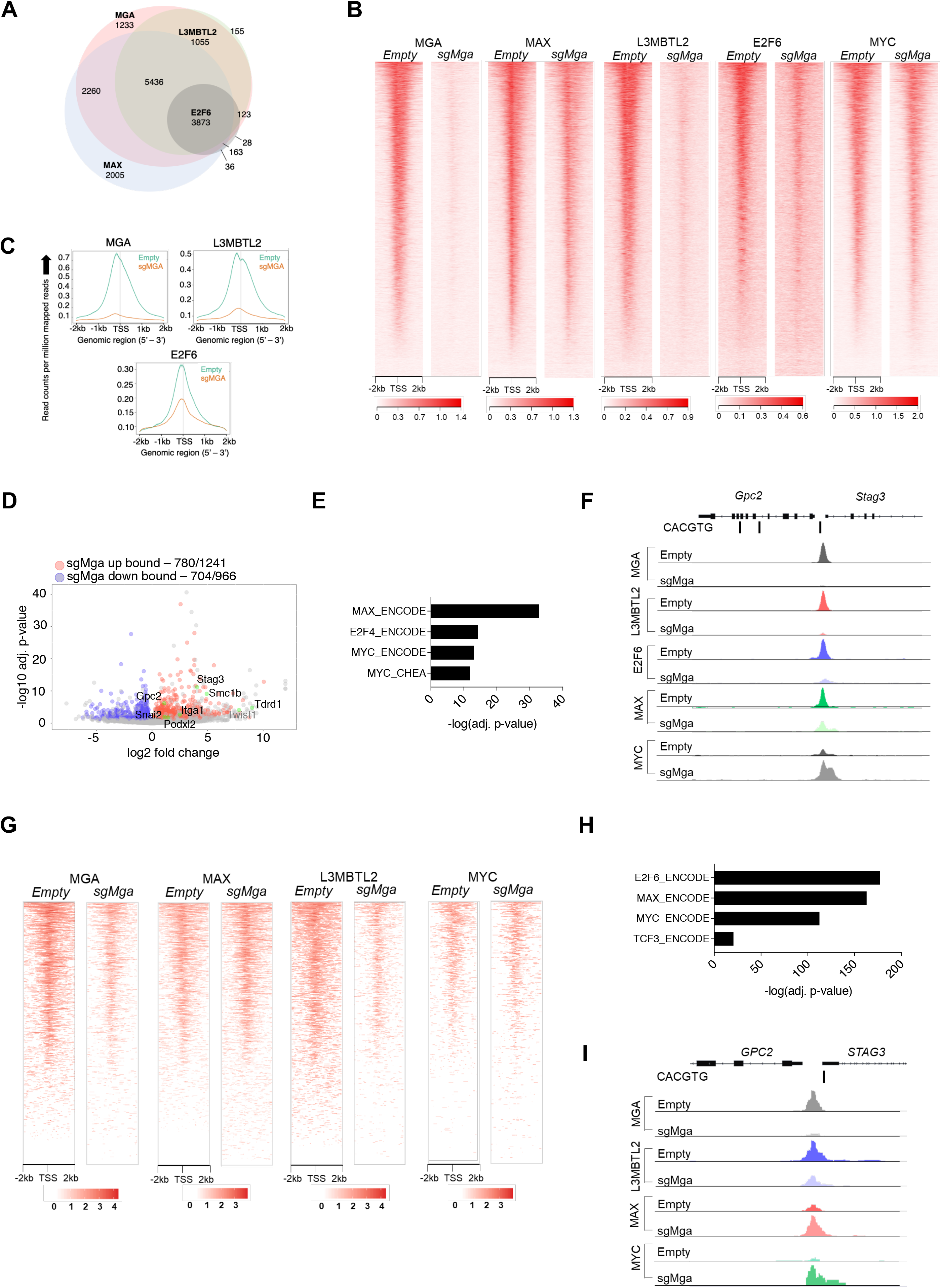
MGA is essential for PRC1.6 genomic binding in tumor cells. (A) Venn diagram showing overlap between MAX, MGA, E2F6 and L3MBTL2 bound genes in KP cells. (B) Heatmaps showing genome wide promoter proximal (+/−2kb) MGA, MAX, L3MBTL2, E2F6 and MYC binding in control and sgMga KP cells. (C) Meta-plots of MGA, L3MBTL2 and E2F6 occupancy in Empty and sgMga KP cells. (D) Volcano plot of differentially expressed genes that are directly bound by MGA (black labels = bound, gray label = unbound). (E) Enrichr analysis of genes bound by MGA (F) representative tracks for MGA. L3MBTL2, E2F6, MAX and MYC binding at the Stag3/Gpc2 promoter. (G) Heatmaps of MGA, MAX, L3MBTL2 and MYC binding in A549 control (Empty) and sgMGA cells. (H) Enrichr analysis of MGA bound genes in A549 cells. (I) Representative tracks for MGA, L3MBTL2, MAX and MYC binding at the STAG3/GPC2 promoter in A549 cells.

To examine whether these results apply to human lung cancer cells, we transduced the MGA expressing A549 human lung cancer line with a guide RNA against MGA and probed for MAX, E2F6, L3MBTL2 and MYC occupancy using CUT&RUN (Janssens et al. 2018). Similar to mouse KP lines, we observed a substantial decrease in MGA and L3MBTL2 binding in *MGA* inactivated cells (termed sgMGA) (Fig. 5G). In addition, Enrichr analysis of MGA bound genes in A549 cells revealed a significant enrichment for MYC, MAX and E2F targets (Fig. 5H). Concomitant with an increase in STAG3 and PODXL2 expression in sgMGA A549 cells, there was decreased occupancy of MGA and L3MBTL2 at their respective promoters, with an increase in MYC binding (Fig. 5I, Fig. S5C). Taken together, this data strongly suggests that MGA mediates PRC1.6 binding to MYC and E2F targets in tumor cells, while loss of MGA abrogates PRC1.6 occupancy at thousands of promoters and leaves MYC-MAX binding either unchanged or increased. Given the substantial enrichment of MYC at PRC1.6 targets such as STAG3/GPC2 and PODXL2, following MGA loss, we wanted to assess the contribution of MYC-MAX heterodimers towards activation of transcription at MGA target genes in MGA deficient cells. First, we examined TCGA data to interrogate co-occurrence of MGA and MYC paralog alterations in a panel of human tumor lines. Several MGA mutated lines amplify MYC paralogs, suggesting that loss of MGA isn’t synthetic lethal with MYC activation (Fig. S5D). Treatment of control and sgMGA A549 cells with 10058-F4, an extensively used MYC-MAX dimerization inhibitor (Yin et al. 2003) revealed that MGA deficient cells are as sensitive to MYC inhibition as Empty transduced cells (Fig. S5E). A similar trend was observed for mouse KP lines upon treatment with another MS2-008 (Struntz et al. 2019), a chemical probe that inhibits MYC-MAX driven transcription (Fig. S5F). Together, these data indicate that MYC is likely to be driving tumor growth in both MGA WT and MGA deleted cells.

### The DUF domain is critical for MGA function

Our genomic occupancy studies confirm that MGA acts as a scaffold for PRC1.6 and that inactivation of MGA results in decreased abundance of PRC1.6 complex subunits and loss of their genomic association. We next wanted to identify the region within MGA that mediates its scaffolding function. Our initial mass spectrometric analysis of immune-precipitates using fragments of MGA in 293FT cells revealed that the region encompassing aa967-1300 is crucial for binding of L3MBTL2 (an obligate component of PRC1.6) and several other proteins (Fig. 4A). To confirm whether this region in MGA is required for binding to L3MBTL2 in vitro we performed co-immunoprecipitation experiments using HA-tagged L3MBTL2 and Flag-tagged MGA or MGAΔ1003-1304 and found clear evidence of preferential association of L3MBTL2 with wildtype MGA as compared with MGAΔ1003-1304 (Fig. 6A). Interrogation of protein domain databases revealed that this region includes a conserved domain of unknown function (DUF4801). Analysis of publicly available sequencing data revealed that mutations occur at a high frequency in this region of MGA (Fig. S6A). To study the functional importance of this region, we generated a mutant version of MGA that lacks these 301aa residues (termed MGAΔDUF). When we ectopically expressed either WT or the DUF mutant version in two different mouse sgMga mouse KP lines, we observed that addback of MGAΔDUF had no effect on proliferation of KP cells whereas expression of WT MGA led to a significant decrease in proliferation (Fig 6B, 6C, 6D). We then stably transduced human MGA mutant lung squamous cell line LOU-NH-91 with FLAG-tagged full-length MGA or MGAΔDUF and determined that both WT MGA and MGAΔDUF forms localize to the nucleus (Fig. S6B). We observed an increase in L3MBTL2 protein levels upon MGA addition but not in the MGAΔDUF expressing cells (Fig. 6E). While there was no effect on MAX levels, MYC levels appeared to decrease in MGA and MGAΔDUF expressing cells, suggesting some form of feedback regulation within the network (Fig. 6E). In contrast to MGA addback, which resulted in a drastic reduction in cell number, MGAΔDUF expression led to a mild impairment in proliferation (Fig. 6F). Concomitantly, we observed a decrease in actin protrusions in MGA overexpressing cells (Fig. 6G). Intriguingly, expression of MGAΔDUF led to an increase in protrusions relative to control cells, suggesting that removal of the DUF domain might result in alterations of gene regulation by MGA (Fig. 6H). This idea is consistent with reports that in Raji leukemia cells, a truncated form of MGA lacking this region activates transcription at E-boxes (Lee et al. 2018). Given that the 301aa region encompassing the DUF domain is critical in recruiting L3MBTL2 and possibly other PRC1.6 components, we assessed whether knockdown of individual PRC1.6 complex members could phenocopy MGA loss. To this end, we used CRISPR to inactivate L3MBTL2 and PCGF6 in A549 cells (Fig. S6C). Neither loss of L3MBTL2 nor PCGF6 phenocopied the effects seen on actin protrusion formation (Fig. S6D) and migration (Fig. S6E, S6F) in MGA CRISPR cells. Overall, this suggests that MGA has functions independent of PRC1.6 and can’t be readily compensated for by other members.

**Figure 6:**
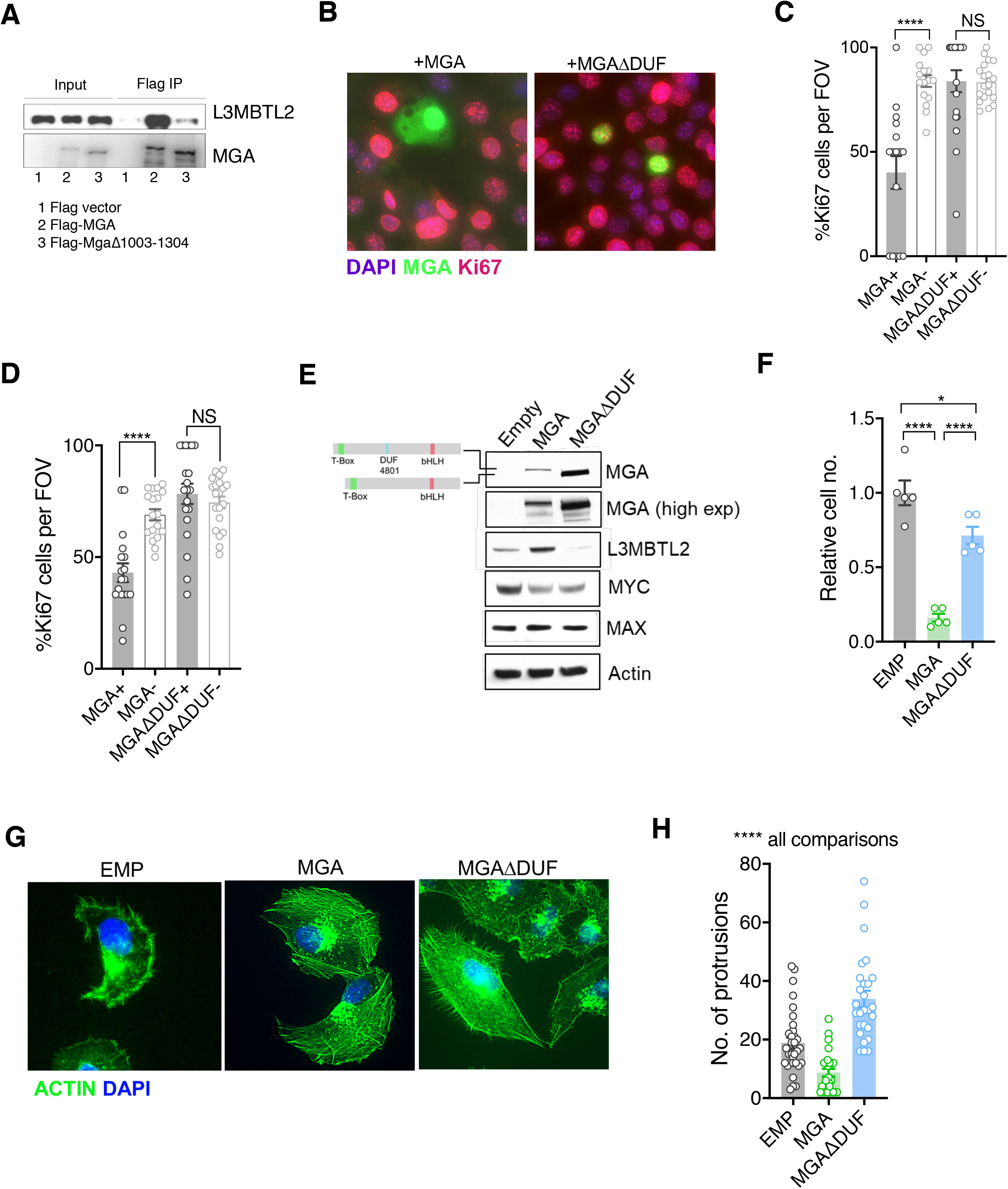
The region containing the DUF4801 domain is critical for MGA’s tumor suppressive function. (A) Immunoblot showing co-immunoprecipitation of L3MBTL2 with full length or MGAΔ1003-1304(DUF). (B) Representative co-staining for Ki67 and MGA deficient mouse KP cells expressing MGA or MGAΔ1003-1304(DUF). Quantification of Ki67+ cells in two independent KP sgMga lines (C) H8712 T3 and (D) H8638. (E) Immunoblots of ectopic expression of MGA or MGAΔDUF in LOU-NH-91 human lung squamous carcinoma cell line. (F) Relative growth of MGA and MGAΔDUF expressing LOU-NH-91 cells. (G) Representative images and, (H) quantification of actin protrusions in MGA, MGAΔDUF expressing and control cells. All p-values calculated using a 2-sided Student’s t-test unless otherwise noted (*p < 0.05, ****p <0.0001). Error bars represent SEM.

### MGA has a tumor suppressive function in colorectal cancer

As mentioned previously, genetic alterations in MGA are prevalent across a broad spectrum of tumors (Schaub et al. 2018). To extend our studies, we aimed to elucidate MGA’s role in other tumor types with a high frequency of MGA alterations. In particular, we were interested in colorectal cancer since we observed a significant percentage of MGA alterations in colorectal adenocarcinoma patients upon interrogation of GENIE and TCGA data (Fig. 7A). In order to study MGA’s role in colorectal cancer initiation, we utilized a CRISPR based approach to inactivate MGA in normal human colon organoids (Fig. 7B) and confirmed that MGA levels were decreased in sgMGA organoids compared to Empty controls (Fig. 7B, Fig. S7A). This was accompanied by a decrease in L3MBTL2 levels, similar to what we observed in lung cancer models (Fig. 7B, Fig. 4B, 4E). We confirmed indel formation by sequencing 2 clonal sgMGA organoids (Fig. S7B). Inactivation of MGA in colon organoids didn’t impact growth in 2D culture (Fig. S7C) but drastically accelerated 3D growth over a period of 14 days (Fig. 7C, 7D,Fig. S7D). To further characterize changes mediated by MGA that facilitate a pro-growth phenotype, we performed global gene expression profiling comparing Empty vector and sgMGA transduced organoids. PCA revealed that sgMGA organoid gene expression profiles form a distinct cluster when compared to Empty controls (Fig. 7E). Importantly, we observed that 4 of the 5 genes (STAG3, PODXL2, NHLRC1 and ZCWPW1) identified from our analysis in lung cancer models were significantly up-regulated upon loss of MGA (Fig. 7F). Enrichment analysis revealed that there are both common and distinct MGA driven transcriptional changes in colon organoids when compared to our murine lung cancer studies. Similar to the mouse lung adenocarcinoma KP model, the expression of several EMT related genes was elevated upon MGA loss. Strikingly, we also noted a broad upregulation of E2F regulated, cell cycle and DNA replication associated genes (Fig. 7G). As in Kras tumors, inflammation and interferon signaling genes were significantly enriched amongst the downregulated genes (Fig. 7H). Several pathways unique to colon organoids also scored significantly. For example, several colon stemness associated genes (van der Flier et al. 2009, Munoz et al. 2012) including LGR5 were upregulated upon MGA loss (Fig. S7E). Conversely, senescence appeared to be a major process enriched amongst the downregulated genes (Fig. 7H, Fig. S7E).

**Figure 7:**
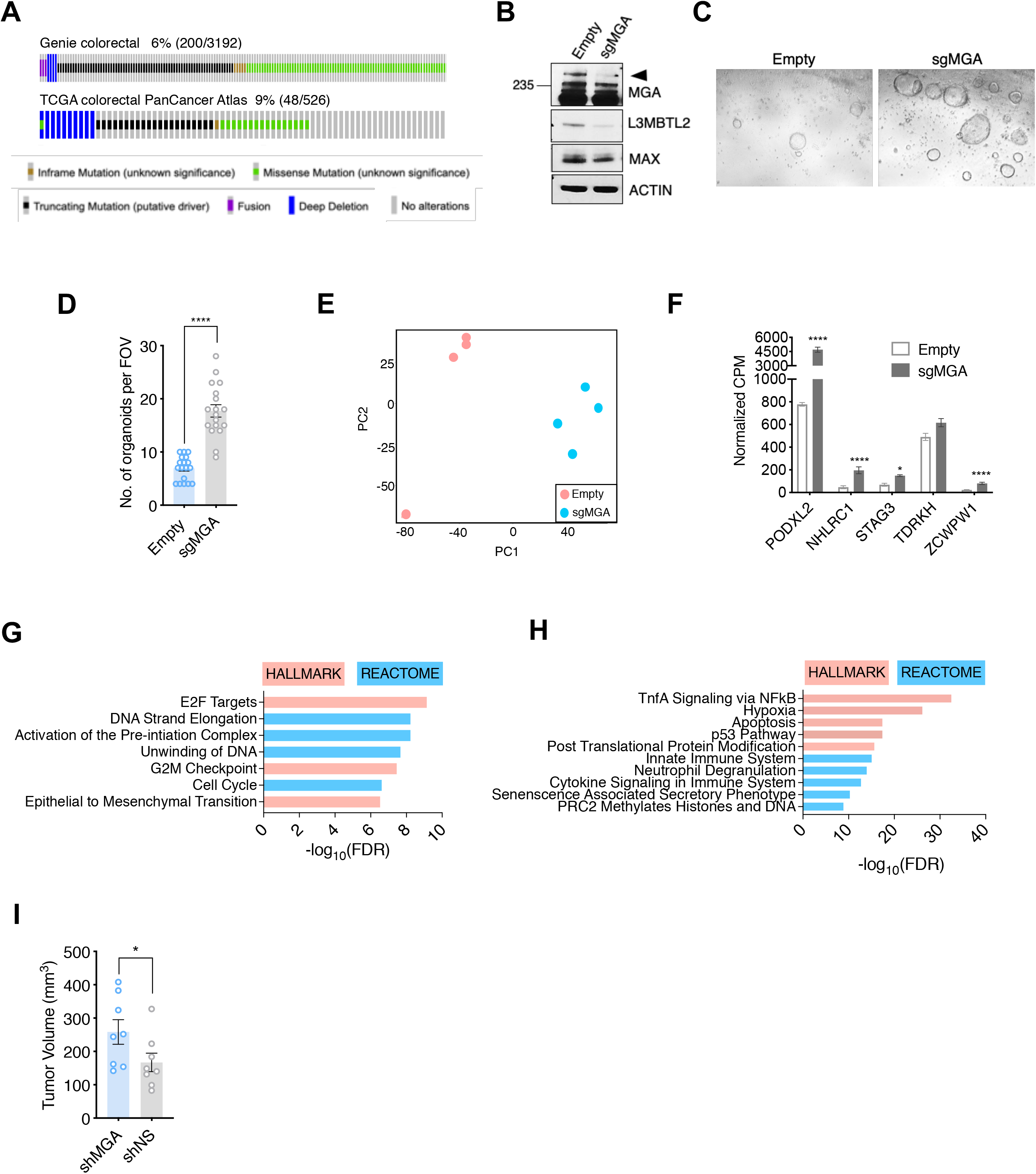
MGA has tumor suppressive functions in colorectal cancer. (A) GENIE and TCGA consortium data depicting alterations at the MGA locus in colorectal cancer. (B) Western blot for MGA, L3MBTL2 and MAX in Empty and sgMGA normal colon organoids. (C) Representative images and, (D) quantification of normal human colon organoid growth following MGA inactivation using CRISPR (*p < 0.0001. 2-sided Students t-test). (E) PCA plots for Empty and sgMGA organoids (n=4 for each). (F) Normalized CPM values for MGA target genes in Empty and sgMGA organoids (****adjusted p-value < 0.0001). GSEA Hallmark and Reactome analysis for genes (G) up-regulated and, (H) down-regulated upon MGA loss in colon organoids (FDR<0.05, LFC>1.5). (I) Tumor volume of subcutaneously implanted shMGA and shNS DLD1 cells 10 days post implantation (n= 8 tumors each. *p < 0.05 calculated using Mann-Whitney test). Error bars represent SEM.

Since the above data establish that MGA loss alters the properties of normal colon cells we next wanted to assess the impact of MGA depletion on malignant colorectal adenocarcinoma cells. We used an shRNA approach to knockdown MGA in DLD1 colorectal adenocarcinoma cells. In agreement with results from lung cancer lines, 2D growth of DLD1 cells was minimally impacted by shRNA knockdown of MGA although shMGA cells exhibited increased actin protrusions (Fig. S7F, S7G). However, shMGA DLD1 xenografts in mice grew faster than non-silencing controls (Fig. 7I). Overall, our data in colorectal cancer models further extend the concept that MGA functions as a tumor suppressor in multiple tissue types.

## DISCUSSION

### Tumor suppressive roles within the extended MYC network in tumorigenesis

MYC family genes are amongst the most intensively studied proto-oncogenes and have been found to be dysregulated across a broad spectrum of malignancies. For example, a recent TCGA pan-cancer analysis found MYC paralogs to be amplified in 28% of all samples (Schaub et al. 2018). MYC proteins belong to a larger network of bHLHZ transcription factors whose members are considered to synergize with or antagonize MYC function (reviewed in Carroll et al. 2018). One subset of network members, the MXD, MNT and MGA proteins act as transcriptional repressors. A role for MNT in antagonizing MYC has been well established (Hurlin et al. 1997, Meroni et al. 1997). *In vivo,* MNT deletion leads to hyperplasia in the mammary gland, similar to MYC overexpression (Toyo-oka et al. 2006), consistent with the idea that MNT acts as a tumor suppressor (Dezfouli et al. 2006). However, MNT has also been shown to collaborate with deregulated MYC in lymphomas by restraining MYC-induced apoptosis (Link et al. 2012, Nguyen et al. 2020). These and other findings suggest that the equilibrium between activating and repressive heterodimers is likely to be critical for tissue homeostasis (Carroll et al. 2018). To investigate the functions of putative MYC antagonists more deeply we focused on MGA, the largest bHLHZ protein known to heterodimerize with MAX. Intriguingly, MGA sustains a high rate of inactivating alterations across a wide range of malignancies (Schaub et al. 2018). We chose to further investigate MGA function in lung cancer since a substantial percentage of lung adenocarcinoma patients (6-10%) harbor MGA alterations (TCGA and GENIE consortium data). Using an *in vivo* CRISPR approach in both Kras and KP models of non-small cell lung cancer, we demonstrate here that MGA loss accelerates tumor progression. Recent studies have suggested that MGA functions as a MYC antagonist and tumor suppressor (Reddy et al. 2017, Llabata et al. 2019), but our work provides strong *in vivo* evidence for its potent tumor suppressive activity. This supports the idea of a broader role for the extended MYC network in lung tumorigenesis, where either activation of MYC paralogs or loss of MYC antagonists results in a pro-malignant phenotype. It is likely that disruption of the balance between MYC-MXD/MNT/MGA antagonism in other tissues can also contribute to tumorigenesis, a concept that warrants further investigation. Indeed, we show that MGA curbs the growth of human colon organoids, strongly suggestive of a tumor suppressive role for MGA in colorectal cancer.

In addition to these systems, MGA may also play a supportive role in tumor suppression under specific circumstances. Several studies indicate that MAX, the obligate heterodimerization partner for both MYC and the MXDs/MNT/MGA, functions as a context specific tumor suppressor in small cell lung cancer and gastrointestinal stromal tumors (Romero et al. 2014, Schaefer et al. 2017, Augert et al. 2020). Small cell lung tumors lacking MAX are independent of MYC but such tumors would also be expected to have inactive MGA as well as MXD/MNT family members, which in principle could contribute to tumorigenesis (Augert et al. 2020).

### MGA functions within the PRC1.6 complex

Recent sequencing efforts reveal that aside from alterations in growth promoting signaling pathways such as in EGFR and KRAS, chromatin modifying enzymes and epigenetic regulators such as MLL3 and TET2 are also frequently altered in malignancies (Kandoth et al. 2013). Dysregulation of polycomb mediated repression through PRC2 is fairly well characterized in several cancers (reviewed in Laugesen et al. 2016). For example, elevated levels of EZH2, a methyltransferase associated with PRC2, drives tumor progression in mouse models of lung adenocarcinoma through the establishment of a unique super-enhancer landscape (Zhang et al. 2016). In addition, studies have established a context dependent function for canonical PRC1 complex members in malignant transformation (Koppens and van Lohuizen, 2016). However, the role of non-canonical PRC1 complexes (ncPRC1) in malignant cells is poorly understood. Recent reports indicate that the four distinct ncPRC1 complexes (PRC1.1; PRC2/4; PRC3/5 and PRC1.6) function in normal development through direct binding to DNA and recruitment of histone modifying enzymes, including methyltransferases, and ubiquitin ligases (Trojer et al. 2011, Gao et al. 2012). PRC1.6 is the only ncPRC1 complex reported to include MAX and MGA. In terms of gene expression, PRC1.6 binding has been associated with permissive transcription and repression (Ogawa et al. 2002, Stielow et al. 2018, Scelfo et al. 2019). In line with these observations in 293T cells, our expression profiling studies in MGA deficient tumors and cell lines revealed a significant up-regulation of known ncPRC1.6 regulated germ cell related transcripts such as *Stag3* and a less marked de-repression of several hundred genes impinging on a diverse array of processes ranging from lipid metabolism to EMT. Importantly, de-repression of certain genes such as STAG3 and PODXL2 was also observed in patient-derived lung adenocarcinoma data (TCGA). We confirmed that these two proteins are critical for the viability of A549 lung cancer cells, underscoring the functional importance of PRC1.6 targets in malignant cells. Our genomic occupancy analyses in both mouse and human lung cancer lines shows that MGA along with L3MBTL2, a defining member of the PRC1.6 complex, directly binds thousands of promoters in tumor cells. Moreover, we observe a substantial overlap of MGA-MAX and L3MBTL2 binding, suggesting that the majority of MGA bound genes in these cells are PRC1.6 regulated. A subset of these genes are also bound by E2F6, consistent with previous studies suggesting that MGA represses at E2F6/ Dp1 sites through PRC1.6 (Ogawa et al. 2002, Llabata et al. 2020). We show that loss of MGA results in greatly diminished L3MBTL2 and E2F6 genomic binding. Taken together with our observation that MGA stabilizes PRC1.6 complex members in tumor cells, our studies strongly implicate MGA as an anchor or scaffold for PRC1.6 even in malignant settings.

### The DUF region is critical for MGA’s tumor suppressive function

Our mass spectrometric analysis using fragments of MGA revealed that aa1000-1300 are critical for MGA interactions with L3MBTL2. This region encompasses a domain of unknown function known as DUF480, and several lines of evidence suggest it is important for mediating tumor suppression. First, addback of a mutant form of MGA lacking this region fails to retard the growth of MGA deficient human and mouse lung cancer lines. Second, several truncating mutations in MGA in the pan-cancer TCGA database appear to be clustered in this region. Indeed, there is a precedent for hotspot mutations in components of chromatin modifying complexes having an oncogenic or tumor suppressive role. For example, mutations frequently occur in the PHD domain of MLL3, a member of the COMPASS complex. These disrupt its binding to the histone H2A deubiquitinase and tumor suppressor BAP1, subsequently leading to reduced BAP1 recruitment to enhancers (Wang et al. 2018). Lastly, MGA truncations in leukemia due to intronic polyadenylation also occur in this region (Lee et al. 2018). The truncated form of MGA produced in this manner acts as a dominant negative and activates MYC and E2F targets. Further experiments will be required to determine if the truncated MGA DUF mutant is oncogenic in certain contexts.

### MGA as a MYC and E2F antagonist in tumors

Numerous studies show that MGA is mutated or deleted across a wide range of malignancies (TCGA 2014, Romero et al. 2014, Reddy et al. 2017). Given the well characterized role of MXDs in repressing MYC targets, it might be expected that MGA exerts a tumor suppressive function via a similar mechanism. Although our study suggests that this is indeed the case, the mode of MGA mediated repression of MYC targets appears to be different from other MXDs. For example, MXD1 and MXI1 form a ternary complex with MAX and mSIN3A/B-HDAC to replace activating MYC-MAX heterodimers, which recruit histone acetylases, during differentiation (Ayer et al. 1995, Schreiber-Agus et al. 1995, Laherty et al. 1997). While we find that a subset of classical MYC targets are upregulated in MGA deficient tumors, what is striking is the large overlap between MYC and PRC1.6 binding in tumor cells to genes that haven’t previously been characterized as MYC targets. A prime example of such a class of genes are meiotic PRC1.6 targets such as STAG3 and TDRD1. Overall, this suggests that during development, in non-transformed cells, MGA and PRC1.6 normally represses inappropriate lineage-specific transcripts (e.g. meiotic genes such as STAG3 as well as pro-invasive genes such as mesenchymal lineage genes like SNAI1). Indeed, we observe a robust upregulation of EMT associated genes and colon stem cell markers upon loss of MGA in normal organoids.

Analysis of our gene expression profiling studies and chromatin occupancy in mouse and human lung cancer models reveal a strong enrichment for E2F4 and E2F6 targets amongst genes bound by and regulated by MGA. In addition, our studies in colon organoids reveal a robust activation of E2F cell cycle gene targets. This is consistent with studies showing that MGA in association with PRC1.6 occupies E2F and MYC responsive genes in resting (G0) cells (Ogawa et al. 2002). A large body of evidence implicates a critical role for MYC and E2F cooperation in maintaining normal tissue homeostasis (Pickering et al. 2009, Liu et al. 2015, Mathsyaraja et al. 2019).

Based on our results, we propose that MGA inactivation in pre-malignant settings destabilizes PRC1.6 and consequentially results in unoccupied E-boxes and E2F/Dp1 sites. A combination of invasion of E-boxes by MYC-MAX and/or binding of E2Fs at targets results in net transcriptional activation of these genes and drives malignant tumor progression (Fig. 8). However, our studies don’t rule out the possibility of other transcription factors contributing to this phenotype. For example, the T-box factor Brachyury is known to promote EMT and invasiveness in several different tumor types, including lung and hepatocellular carcinoma (Fernando et al. 2010, Du et al. 2014, Shah et al. 2017, reviewed in Chen et al. 2020). It is conceivable that MGA mediates repression of EMT related transcripts via its TBX domain. Further structure-function studies will enable the delineation of the role of MGA’s T-box domain in suppressing tumorigenesis.

**Figure 8.**
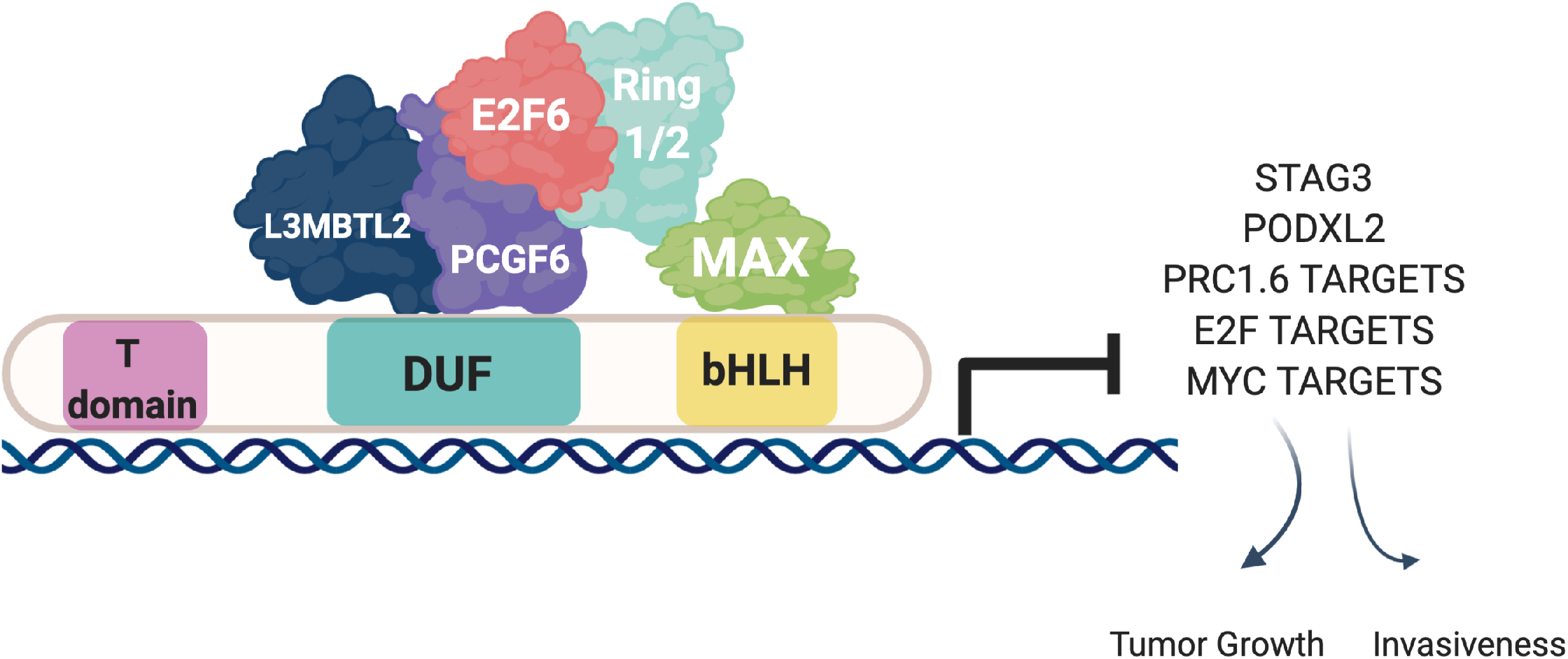
Proposed mechanism of MGA mediated tumor suppressive effects: MGA acts as a scaffold and stabilizes atypical PRC1.6 members, including L3MBTL2. Under normal conditions, this results in the repression or transcriptional attenuation of thousands of genes. During malignant progression, perturbation of MGA expression leads to the up-regulation of growth promoting and pro-invasive PRC1.6, MYC and E2F targets in a tissue specific manner.

### MGA and its potential role in promoting anti-tumor inflammation

Our gene expression profiling data reveal a surprising link between MGA and tumor inflammation. In mouse Kras tumors, human TCGA data and colon organoids, loss of MGA expression correlates with decreased expression of genes involved in inflammatory and interferon gamma responses. It is conceivable that MGA is involved in invoking and maintaining an anti-tumor inflammatory response which contributes to its tumor suppressive function. Downregulation of these anti-tumor inflammation genes upon MGA loss may reflect a transcriptional activation function associated with MGA and ncPRC1.6 (Scelfo et al. 2019). Indeed, it was earlier reported that MGA can act as an activator at E-boxes and T-box binding motifs (Hurlin et al. 1999). Alternatively, MGA may antagonize the binding of MYC-MIZ complexes that have been reported to directly repress these cadre of genes (Muthalagu et al. 2020, Swaminathan et al. 2020). A third possibility is that MGA plays an indirect role in activating interferon pathway genes. Studies in A549 lung cancer cells reveal that MGA acts as host restriction factor in which decreased MGA expression is associated with increased susceptibility to infection by several animal viruses including Sindbis and influenza viruses (Varble et al. 2013). The possibility of MGA acting as a transcriptional activator in tumors remains to be tested and further studies will be required to delineate the specific components of the immune microenvironment that are impacted by MGA loss in human malignancies.

In summary, we demonstrate that MGA acts as a bona fide tumor suppressor *in vivo* and have uncovered a critical role for MGA and the atypical PRC1.6 complex in lung adenocarcinoma. We also provide preliminary evidence for a tumor suppressive role for MGA in colorectal cancer. Although our data is suggestive of a broad tumor suppressive role of MGA and PRC1.6 via antagonizing transcription of MYC and E2F targets, further studies will be required to elucidate the functional relevance of individual MGA/PRC1.6 targets such as STAG3 in tumorigenesis. Given the frequent inactivation of MGA across diverse cancer types, it will also be critical to further characterize MGA and PRC1.6 mediated transcriptional attenuation in other malignancies with MGA alterations, such as colorectal cancer and diffuse large B cell lymphoma. This is especially important as MGA, via atypical PRC1.6, likely regulates lineage specific genes to mediate its tumor suppressor role in a context dependent manner.

## MATERIALS AND METHODS

### Mouse models and generation of cell lines

All mice used in the study were housed and treated according to the guidelines provided by the Fred Hutch Institutional Animal Care and Use Committee. The Kras G12D LSL and Trp53 floxed alleles were maintained on a C57BL6 background. Heterozygous Kras G12D LSL/+ alone (termed Kras) or in combination with Trp53 fl/fl homozygous alleles (termed KP) were used for experiments. Mice 2 months or older were utilized for intratracheal instillation of lentiviral particles in a BSL2 facility (DuPage et al. 2009). For generation of KP mouse lines, tumors were collected using sterile surgical instruments from endpoint KP mice. Tumors were mechanically disaggregated using a wide bore pipette tip and cultured in DMEM supplemented with 10% FBS and antibiotics. Cells were passaged until tumor cells outgrew stromal contaminants (immune cells, fibroblasts and endothelial cells) and used for subsequent downstream applications.

### Cloning, transfection and lentiviral transduction

For in vivo CRISPR, two sgRNA targeting mouse Mga (sgMga #1 and sgMga#3) were designed and cloned into the lentiCRISPRv2cre vector (Walter et al. 2017). See Key Resources table for sequences. Virus was generated by the Fred Hutch Cooperative Center for Excellence in Haematology Viral Vector core and titered using qPCR. For in vitro studies in human lung cancer lines, a guide RNA against MGA was cloned into the lentiCRISPRv2 puro vector (kind gift from Feng Zhang). Viral supernatant was generated via transfection of 293FT or 293TN cells with lentiCRISPR v2 constructs and packaging vectors pPAX2 and pVSV-G using lipofectamine 2000 (Invitrogen). Supernatant was cleared of cellular debris using a 0.45uM filter and target cells were transduced with viral supernatant containing 4ug/ml polybrene final concentration. Cell were grown in 10ug/ml puromycin 3 days post transduction. For addback of MGA or MGAΔDUF to mouse KP lines, cells were transfected on chamber slides with pCDH-MGA-FLAG or pCDH-MGAΔDUF-FLAG using lipofectamine 2000 (Invitrogen). Cells were harvested 3 days post transfection and fixed for immunofluorescent staining.

### Cell lines and reagents

Mouse KP and KP-sgMga lung tumor cell lines were generated as described above and cultured in DMEM with 10% FBS. Human non-small cell lung cancer lines NCI-H23, NCI-H2291, NCI-H2347, NCI-H1975 and 91T were grown in RPMI-1640 with 10% FBS. Human squamous cell carcinoma line LOU-NH-91 was maintained in RPMI-1640 supplemented with 20% FBS. A549 human lung adenocarcinoma, DLD1 human colorectal adenocarcinoma, 293FT and 293TN cells were maintained in DMEM with 10% FBS.

### Human colon organoid culture and maintenance

Organoid cultures were established as previously described (Guo et al. 2019). Briefly, organoids were plated in Matrigel (Corning #354230) and IntestiCult organoid growth media (StemCell #6010) was supplemented with 10uM of ROCK Inhibitor Y27632 (ATCC ACS-3030) for one-week post establishment. For lentiviral transduction, supernatant was added to individual wells of cells in ultra-low attachment plates. The plate was centrifuged at 600g at 32°C for 1 hour. Plate was removed from centrifuge and incubated for 37°C for 4 hours. Cells were washed in cold 1x PBS containing 10uM of ROCK Inhibitor Y27632 and pelleted. Cells were then plated in a 10uL ring of Matrigel in a 96-well tissue culture plate. After Matrigel hardened, 100uL of IntestiCult containing 10uM of ROCK Inhibitor Y27632 was added to each well. Two days post plating 4ug/ml of puromycin (Sigma #P8833) was added to culture media and refreshed every 48 hours. Single cell organoid colonies were established via single cell dissociation using TrypLE Express enzyme, described above. Single cells were plated in 10ul Matrigel rings in a 96-well plate and expanded. Organoid viability was measured using RealTime-Glo MT Cell Viability Assay (Promega #G9711), following manufacturer’s protocol. Single cell organoids were plated at a density of 5×10^3^ cells per well in a 96-well, 2-D culture format, described previously (Thorne et al. 2018). Bright field images were acquired using Nikon Eclipse E800 fluorescent microscope.

### DNA extraction and Sequencing for indels

To confirm indels, cell pellets and tumors were incubated in a Proteinase K containing digestion buffer overnight at 55C. DNA was isolated using either 25:24:1 Phenol:chloroform:isoamylalcohol followed by ethanol precipitation or extracted using the GeneJet genomic DNA purification kit (ThermoFisher). For human colon organoids, cells were dissociated into single cells and DNA was isolated using the Purelink Genomic DNA isolation kit (Invitrogen). PCR was performed using high fidelity Phusion DNA polymerase (ThermoFisher) with primers flanking the region targeted by sg*Mga*#1 and sg*Mga*#3 (for mouse tumors and cell lines) and sgMGA (for human colon organoids). PCR products obtained were run on a 0.8% agarose gel and amplicons to confirm the presence of one amplicon corresponding to the correct size. PCR DNA was purified using a PCR purification kit (Qiagen) and sequenced using one of the sg*Mga#1* sg*Mga#3* (mouse) and sgMGA (human) primers. Sequence information was input for ICE analysis (Synthego) to compute percent indel formation (https://ice.synthego.com) and Clustal Omega (https://www.ebi.ac.uk/Tools/msa/clustalo/) was used to align sequences.

### Immunostaining and Western blots

Cells were cultured on 8-well chamber slides for immunocytochemistry. Briefly, cells were fixed in either 4% formaldehyde or 1:1 methanol:acetone, blocked and incubated with primary antibodies followed by Alexa fluor conjugated secondary antibodies or Alexa 488 conjugated phalloidin (for Actin staining). Chambers were then removed and slides mounted using Prolong Gold anti-fade with DAPI. Immunohistochemistry was performed on 5um thick paraffin embedded mouse lung sections. Following heat-induced antigen retrieval, sections were incubated with primary antibodies followed by Alexa conjugated secondary antibodies. Sections were mounted in Prolong Gold anti-fade with DAPI (Invitrogen). Images were acquired using either TissueFaxs (Tissugnostics), Nikon E800, Deltavision Eclipse, Olympus Fluoview confocal or Zeiss confocal microscopes. Image analysis and intensity measurements were performed using ImageJ.

For protein gels, whole cell lysates were prepared using RIPA buffer with protease and phosphatase inhibitors. They were then reduced in NuPage LDS buffer (Invitrogen) and a wet transfer was performed prior to western blotting. For histone blots, acid extracts were made following manufacturer (Abcam) recommendations. Refer to Key Resource Table for list of antibodies used.

### RNA isolation, qPCR and RNA-Sequencing

For all applications, RNA was isolated using Trizol (Thermo Fisher Scientific) according to manufacturer’s recommendations or processed using the Direct-zol RNA Miniprep kit (Zymoresearch). For qPCR, cDNA was synthesized from 500ng-2μg of total RNA using the Revertaid cDNA synthesis kit (Thermo-Fisher). A Biorad iCycler was using for SYBR green based quantitative PCR. Refer to supplemental tables 1 and 2 for primer sequences. For RNA sequencing experiments, Kras frozen tumors or KP cell lines were used. Following RNA isolation, total RNA integrity was checked using an Agilent 4200 TapeStation and quantified using a Trinean DropSense96 spectrophotometer (Caliper Life Sciences). Libraries prepared using the either the TruSeq RNA Sample Prep v2 kit (Illumina) with 500ng input RNA or Ultra RNA Library Prep Kit for Illumina (New England Biolabs). An Illumina HiSeq 2500 was utilized to performed paired-end sequencing.

Reads that didn’t pass Illumina’s base call quality threshold were removed and then aligned to mm10 mouse reference genome using TopHat v2.1.0. Counts were generated for each gene using htseq-count v0.6.1p1 (using the “intersection-strict” overlapping mode). Genes that didn’t have at least 1 count/million in at least 3 samples were removed. Data was normalized and comparisons conducted using the exact test method in edgeR v3.18.1. Gene set enrichment analysis was performed using Hallmark and Reactome datasets on mSigDB (Subramanian et al. 2005, Liberzon et al. 2015). Heatmaps were generated using Morpheus (https://software.broadinstitute.org/morpheus).

### Tandem affinity purification of MGA-interacting proteins and mass spectrometry

Twenty 15-cm plates of 293T cells were transfected with pcDNA3-FLAG-His-MGA, MGA(967 – 1300), or MGA(2153 – 2856); 48 hours after transfection, the cells were lysed in TN buffer (10 mM Tris pH 7.4/150 mM NaCl/1% NP-40/1 mM AEBSF/10 μg/ml aprotinin/10 μg/ml Leupeptin/1 μg/ml Pepstatin A/20 mM sodium fluoride). The lysate was incubated with Ni-NTA agarose (Qiagen) and FLAG-His-MGA and its interacting proteins were collected by centrifugation, washed three times with TN buffer, and eluted with 50 mM sodium phosphate buffer pH 8.0/150 mM NaCl/250 mM imidazole. The eluted sample was immunoprecipitated with anti-FLAG antibody (M2, Sigma-Aldrich), the immunoprecipitate was eluted with FLAG peptide (Sigma-Aldrich), and the eluted protein sample was processed with an Amicon Ultra 0.5 3k centrifugal filter device (Millipore) for concentration and buffer exchange to 50 mM Tris pH 8.5. Proteins were digested at 37 °C overnight with trypsin (Promega; 1:10, enzyme/substrate) in the presence of 10% acetonitrile. The resulting tryptic peptides were analyzed by HPLC-ESI-tandem mass spectrometry on a Thermo Fisher LTQ Orbitrap Velos Pro mass spectrometer. The Xcalibur raw files were converted to mzXML format and were searched against the UniProtKB/Swiss-Prot human protein database (UniProt release 2016_04) using X! TANDEM CYCLONE TPP (2011.12.01.1 - LabKey, Insilicos, ISB). Methionine oxidation was considered as a variable modification in all searches. Up to one missed tryptic cleavage was allowed. The X! Tandem search results were analyzed by the Trans-Proteomic Pipeline, version 4.3. Peptide/protein identifications were validated by the Peptide/ProteinProphet software tools (Keller et al. 2002; Nesvizhskii et al. 2003).

### Genomic occupancy analyses

For genomic occupancy studies in mouse KP lines, conventional crosslinked ChIP-seq was performed (Skene and Henikoff. 2015) on one *Mga* inactivated line and one control line. Chromatin IPs were done using antibodies against MGA, MAX, MYC, L3MBTL2 and E2F6. Libraries were generated using the NEB Ultra II kit (E7645S, NEB), followed by 50X50 paired end sequencing on an Illumina HiSeq 2500 instrument. For human cell lines, CUT&RUN was utilized to determine genomic occupancy. Cells were bound to Concanavalin A beads (86057-3, Polysciences Inc), followed by permeabilization using a digitonin containing buffer. 1 million cell aliquots were then incubated overnight with antibodies against MGA, MAX, MYC and L3MBTL2. An automated CUT& RUN protocol was followed for library preparation (Janssens et al. 2018) and libraries were sequenced using 25×25 paired-end sequencing on an Illumina HiSeq 2500 instrument. 5-10 million reads were obtained per antibody. Refer to Key Resource table for list of antibodies used.

For ChIP-seq experiments, sequences were aligned to the mm10 reference genome assembly using Bowtie2. For CUT&RUN experiments, alignment was performed to hg38. Library normalization was utilized for ChIP-Seq experiments and spike-in normalization was performed using yeast DNA spike-in for CUT&RUN studies. Peak calling was performed using MACS at different thresholds. Peaks called were further processed using bedtools plus a combination of custom R scripts defining genome position and the GenomicRanges R package. For MGA, peaks were identified as being associated with a gene if they were within + or − 5kb from the TSS. For CUT&RUN experiments, the intersection of MGA peak calls from two independent experiments following IgG subtraction was utilized to obtain a gene list. The R package ggplot2 or ngs.plot (Shen et al. 2014) were used to generate heatmaps for genomic binding. Volcano plots were generated using the R package ggplot2. Enrichr was utilized to overlap peak calls with existing ENCODE and ChEA data. De novo and known motif enrichment for sequence specificity was determined using HOMER (Heinz et al. 2010).

### Cell growth, spheroid and migration assays

For two dimensional growth curves, cells were seeded in 96 well flat bottom dishes (Corning) and imaged at fixed time intervals using either an Incucyte S3 or Incucyte zoom (Essen Bioscience). Percent confluence was used as a measure of cell growth. For 3D spheroid based assays, 5000 A549 cells were plated per well in ultra-low attachment plates and centrifuged at low speed (800rpm) for 9 minutes. Phase contrast images were acquired using an inverted scope. Spheroid area was calculated using ImageJ. For wound healing assays, cells were plated to reach full confluence prior to the start of the assay. Imagelock plates (Essen Bioscience) were utilized. Cells were pre-treated with mitomycin C (2.5-5 ug/ml) to retard cell growth and scratch wounds were made using a Woundmaker (Essen Bioscience). Wound closure was monitored at fixed time intervals and endpoint wound width or confluence were used as metrics for cell migration.

## ACKNOWLEDGEMENTS

We would like to thank the members of the Eisenman and MacPherson labs for scientific input and sharing reagents. We are grateful to Guntram Suske, Bastian Stielow, Akihiko Okuda and Peter Hurlin for their generous gifts of key reagents, to Alice Berger for helpful discussions and to Arnaud Augert for a critical reading of the manuscript. We also acknowledge the Fred Hutch Genomics and Bioinformatics, Scientific Imaging, Experimental Histopathology, and Small Animal Imaging Shared Resources for their excellent technical support. We are also grateful for technical help provided by the Cooperative Center for Excellence in Haematology (CCEH) at Fred Hutch. We thank the University of Texas Health Science Center at San Antonio Institutional Mass Spectrometry Laboratory (Director: Dr. Susan Weintraub) for mass spectrometry analysis (supported by P30CA054174). This work was supported in part by NIH/NCI grants R35CA231989 (to R.N.E.), U01CA235625 and R01CA200547 (to D.M.), and by a Hartwell Innovation Fund Pilot Grant (to R.N.E.). This work was also funded in part by NIH/NCI (CA202485), Cancer Prevention and Research Institute of Texas (RP160487, RP160841, and RP190385), the Owens Medical Research Foundation (to Y.S); NIH/NCI R50CA233042 (to M.Y.), UO1CA152756, RO1CA220004, P30CA015704, Brotman Baty Institute Pilot award, Cottrell Family Fund, Geiger Family Foundation and Listwin Fund (to W.M.G.).

