## Supplemental figures+information for "Loss of MGA mediated Polycomb repression promotes tumor progression and invasiveness"

### Figure S1

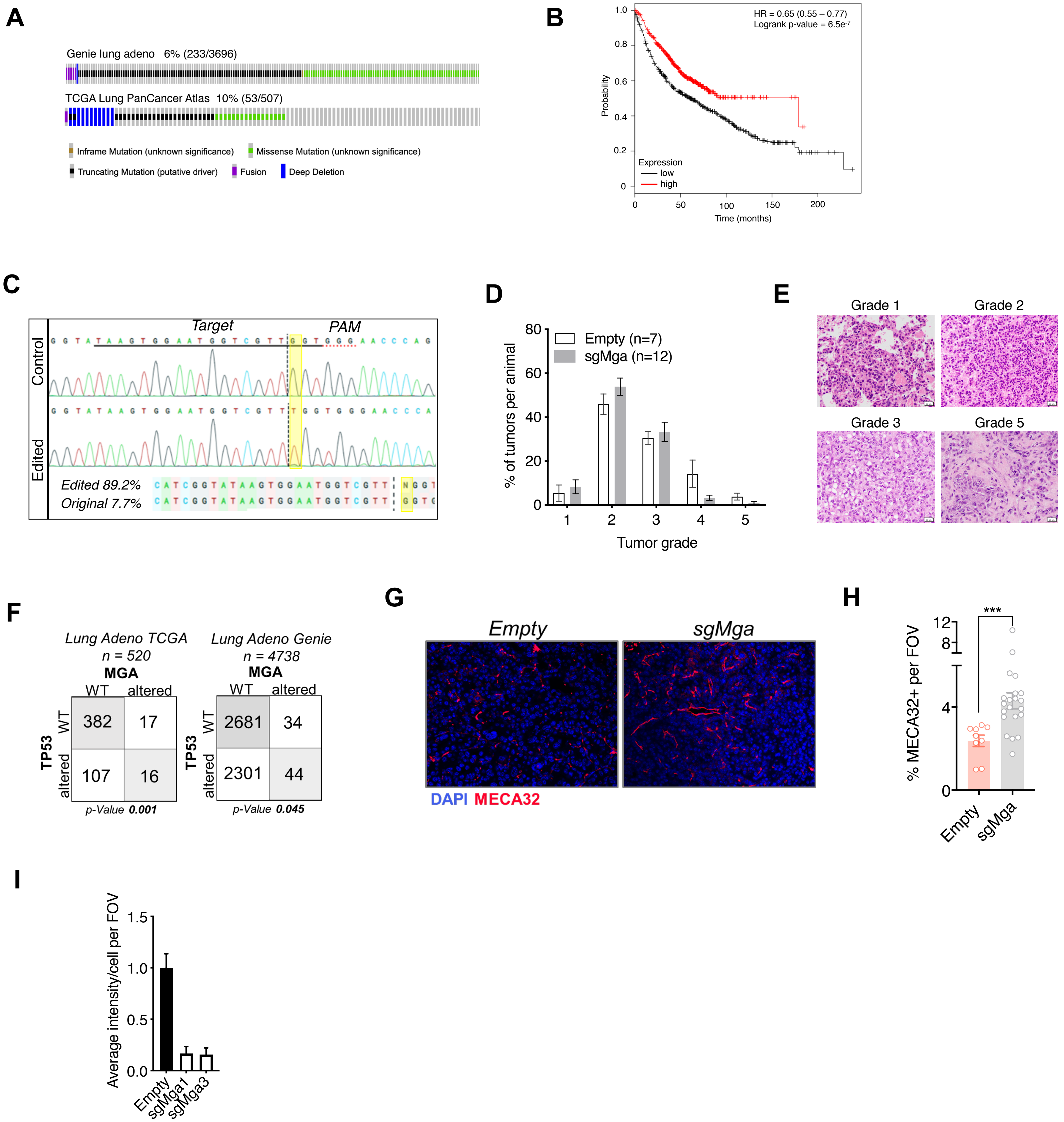

A
Genie lung adeno 6% (233/3696)
TCGA Lung PanCancer Atlas 10% (53/507)
Inframe Mutation (unknown significance)
Missense Mutation (unknown significance)
Truncating Mutation (putative driver)
Fusion
Deep Deletion
B
HR = 0.65 (0.55 – 0.77)
Logrank p-value = 6.5e-7
Probability
Expression
low
high
Time (months)
C
Control
Edited
Target
PAM
Edited 89.2%
Original 7.7%
D
Empty (n=7)
sgMga (n=12)
% of tumors per animal
Tumor grade
E
Grade 1
Grade 2
Grade 3
Grade 5
F
Lung Adeno TCGA
n = 520
MGA
WT altered
TP53
WT 382 17
altered 107 16
p-Value 0.001
Lung Adeno Genie
n = 4738
MGA
WT altered
TP53
WT 2681 34
altered 2301 44
p-Value 0.045
G
Empty
sgMga
DAPI MECA32
H
% MECA32+ per FOV
***
Empty
sgMga
I
Average intensity/cell per FOV
Empty
sgMga1
sgMga3

**Figure S1: *Mga* inactivation *in vivo* leads to accelerated tumorigenesis. (**A) Oncoprint of TCGA and GENIE consortium data on genetic alterations in MGA in lung adenocarcinoma patients. (B) Kaplan-Meier curve for survival probability based on low and high expression of MGA in lung adenocarcinoma patients (KM plotter). (C) Synthego ICE analysis of representative tumor to show percentage of indels formed upon Mga CRISPR in tumors. Example shows a frameshift mutation (resulting from insertion of an additional Thymidine) at a frequency of 89.2%. (D) Percentage of tumors of each grade in Empty vs. sgMga infected Kras mice harvested at endpoint. (E) Representative H&E images of tumor grades seen in Kras mice. (F) Co-occurrence of MGA and TP53 alterations in TCGA and GENIE consortia data from lung adenocarcinoma patients. (G) Representative micrographs and, (H) quantification of pan endothelial marker MECA32 in Empty and sgMga treated KP mice harvested 3 months post intratracheal instillation (Empty: n=9 tumors from 3 mice, sgMga: n=21 tumors from 2 mice. ***$p<0.001$ using Welch's t-test assuming unequal variance). (I) Quantification of MGA intensity in Empty, sgMga1 and sgMga3 KP cell lines (n=5 FOV for all groups). Error bars represent SEM.

### Figure S2

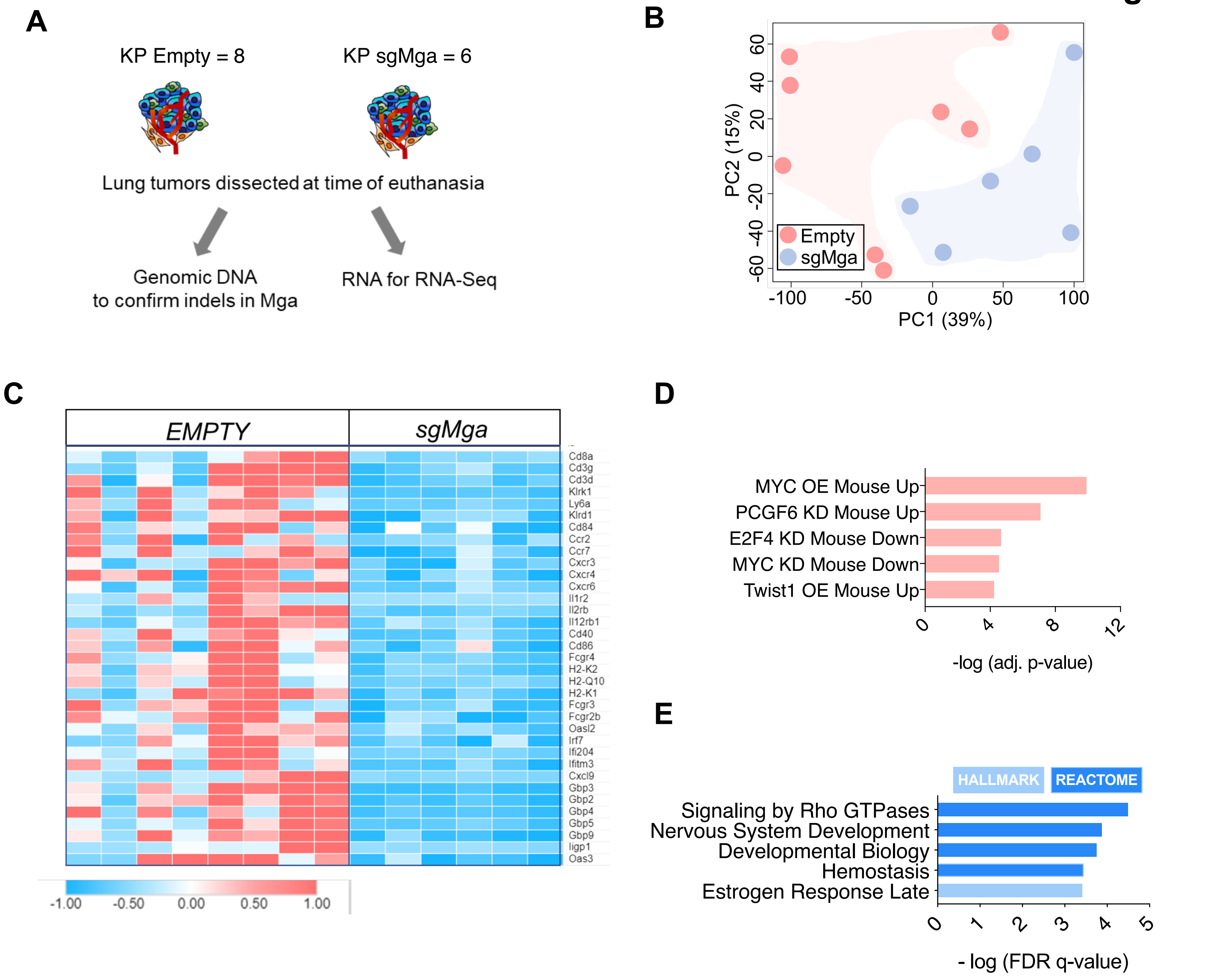


**Figure S2: MGA loss leads to the de-repression of PRC1.6 and MYC targets and up-regulation of pro-invasion genes.** (A) Schematic depicting workflow of Kras tumor RNA Seq. (B) PCA analysis of sgMGA and Empty RNA Seq (n=8 Empty n=6 sgMga). (C) Heatmap of downregulated immune cell and inflammation related transcripts in Kras sgMga tumors compared to Empty. (D) Transcription Factor Perturbation datasets enriched in sgMGA KP upregulated genes (using Enrichr). (E) GSEA analysis of genes downregulated upon loss of MGA in KP lines.

### Figure S3

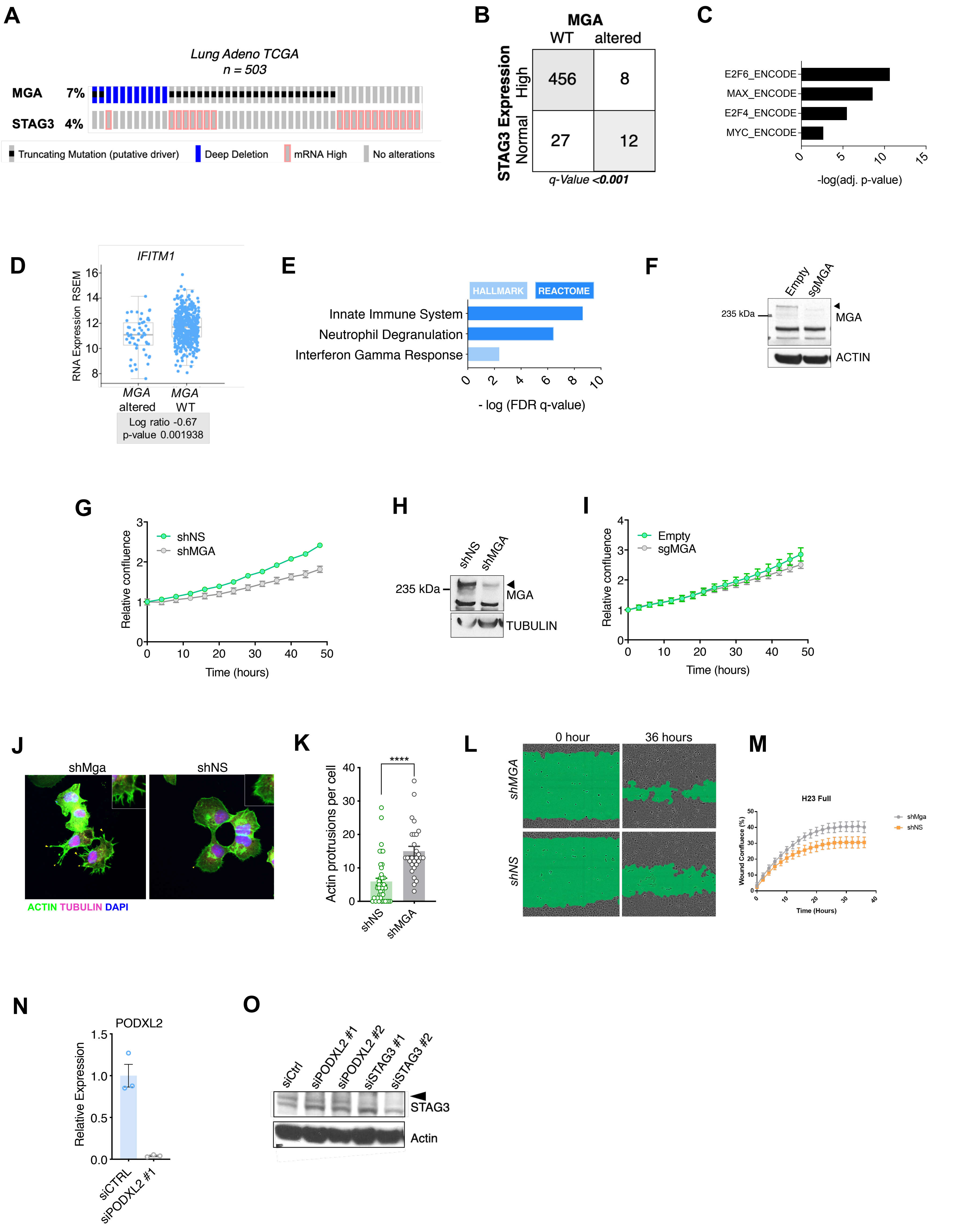

A
Lung Adeno TCGA
n = 503
MGA 7%
STAG3 4%
Truncating Mutation (putative driver)
Deep Deletion
mRNA High
No alterations
B
MGA
WT
altered
STAG3 Expression
High
Normal
456
8
27
12
q-Value <0.001
C
E2F6_ENCODE
MAX_ENCODE
E2F4_ENCODE
MYC_ENCODE
-log(adj. p-value)
D
IFITM1
RNA Expression RSEM
MGA altered
MGA WT
Log ratio -0.67
p-value 0.001938
E
HALLMARK
REACTOME
Innate Immune System
Neutrophil Degranulation
Interferon Gamma Response
- log (FDR q-value)
F
Empty
sgMGA
235 kDa
MGA
ACTIN
G
shNS
shMGA
Relative confluence
Time (hours)
H
shNS
shMGA
235 kDa
MGA
TUBULIN
I
Empty
sgMGA
Relative confluence
Time (hours)
J
shMga
shNS
ACTIN TUBULIN DAPI
K
****
Actin protrusions per cell
shNS
shMGA
L
0 hour
36 hours
shMGA
shNS
M
H23 Full
shMga
shNS
Wound Confluece (%)
Time (Hours)
N
PODXL2
Relative Expression
siCTRL
siPODXL2 #1
O
siCtrl
siPODXL2 #1
siPODXL2 #2
siSTAG3 #1
siSTAG3 #2
STAG3
Actin

**Figure S3: Manipulation of MGA in human lung cancer cell lines results in a pro-invasive phenotype and activation of PRC1.6 targets.** (A) Genetic alteration in MGA and expression changes in STAG3 in human lung adenocarcinoma. (B) Co-occurrence of STAG3 ‘mRNA high expression’ and MGA alterations in TCGA lung adenocarcinoma data. (C) ChEA analysis of genes overexpressed in MGA altered cases vs. WT (D) Expression of IFN signaling gene IFITM1 in MGA altered vs. WT lung adenocarcinoma patients. (E) GSEA on genes underexpressed in MYC amplified vs. non-amplified cases. (F) Immunoblot of MGA levels in sgMGA and Empty vector transduced A549 human lung adenocarcinoma cells. (G) Growth curve of Empty and sgMGA A549 cells cultured in complete media. (H) Immunoblot of MGA levels in control non-silencing shRNA (shNS) and shMGA NCI-H23 lung adenocarcinoma cells. (I) Growth curve of shMGA and non-silencing control NCI-H23 cells. (J) Representative micrograph and (K) quantification of phalloidin (Actin) staining of shNS and shMGA H23 cells (****$p<0.0001$) (L) Wound width at 36 hours and (M) kinetics of wound closure in shNS and shMGA transduced NCI-H23 cells. (N) qPCR for PODXL2 in siCTRL vs. siPODXL2#1 transfected A549 Empty cells (n=3 for each condition). (O) Western blot for STAG3 levels in siCTRL, siPODXL2 and siSTAG3 transfected A549 empty cells. All p-values calculated using a 2-sided Student’s t-test unless otherwise noted. Error bars represent SEM.

### Figure S4

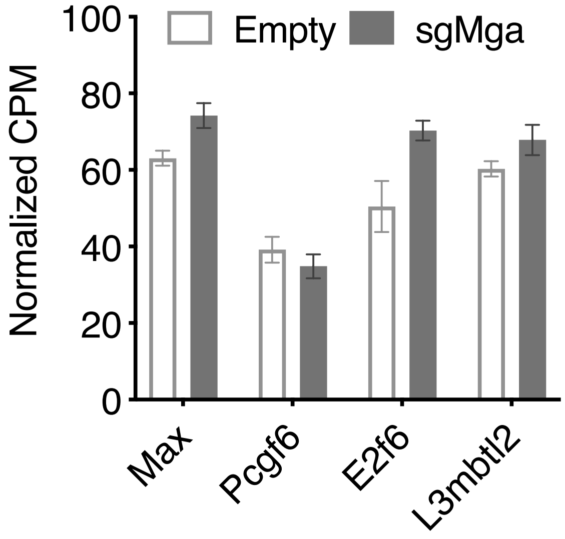


**Figure S4: MGA stabilizes PRC1.6 complex members in lung cancer cells.** Normalized CPM values for PRC1.6 member expression in KP cells (n=2 Empty lines in triplicate, n=3 sgMga lines in triplicate). Error bars represent SEM.

### Figure S5

A B C

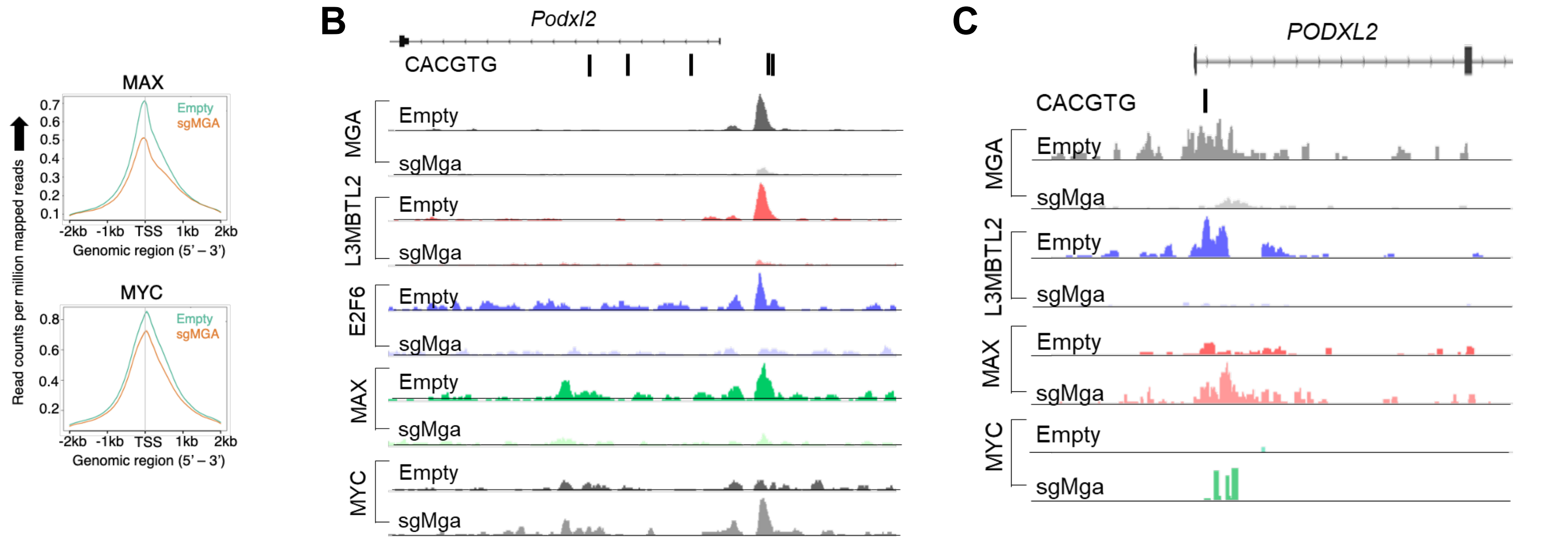


D

| Cell line | Tumor Type | MYC amplification | MGA alteration |
|---|---|---|---|
| 59M | Ovarian | Yes | Deletion |
| MSTO-211H | Mesothelioma | Yes | Deletion |
| OVCAR-8 | Ovarian | Yes | Deletion |
| MDA-MB-453 | Breast | Yes | Mutation S697* |
| 23132_87 | Stomach | Yes | Mutation R896* |
| DMS153 | SCLC | Yes (MYCL) | Deletion |
| COR-L47 | SCLC | Yes (MYCL) | Mutation E2035* |
| LXF-289 | NSCLC | Yes (MYCL) | Mutation Q1067* |

E

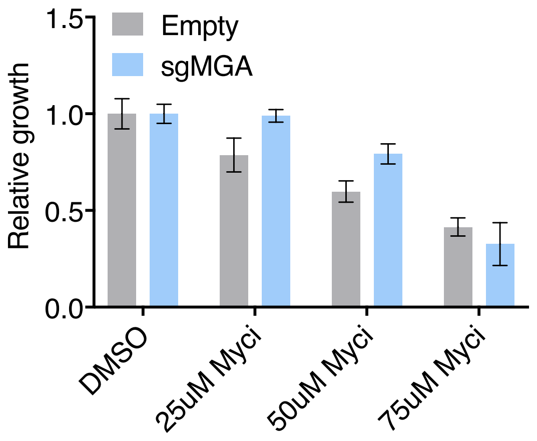


F

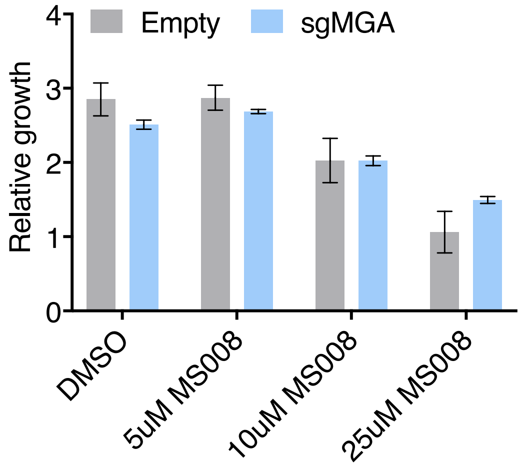


**Figure S5: MGA is essential for PRC1.6 genomic binding in tumor cells** (A) Line plots of IgG, MAX, and MYC enrichment at the TSS +/- 2kb in control and sgMGA KP lines. (B) Peaks for MGA, L3MBTL2, E2F6, MAX and MYC occupancy at the *Podxl2* promoter in control KP cells. (C) MGA, L3MBTL2, MAX and MYC occupancy at the *PODXL2* promoter in control A549 cells. (D) Table showing co-occurrence of alterations in MGA and MYC paralogs in cancer cell lines. (E) Relative growth of A549 Empty and sgMGA cells in varying concentrations of Myci( 10058-F4). (F) Viability of A549 lines (Empty and sgMga) treated with MS2-008. Error bars represent SEM.

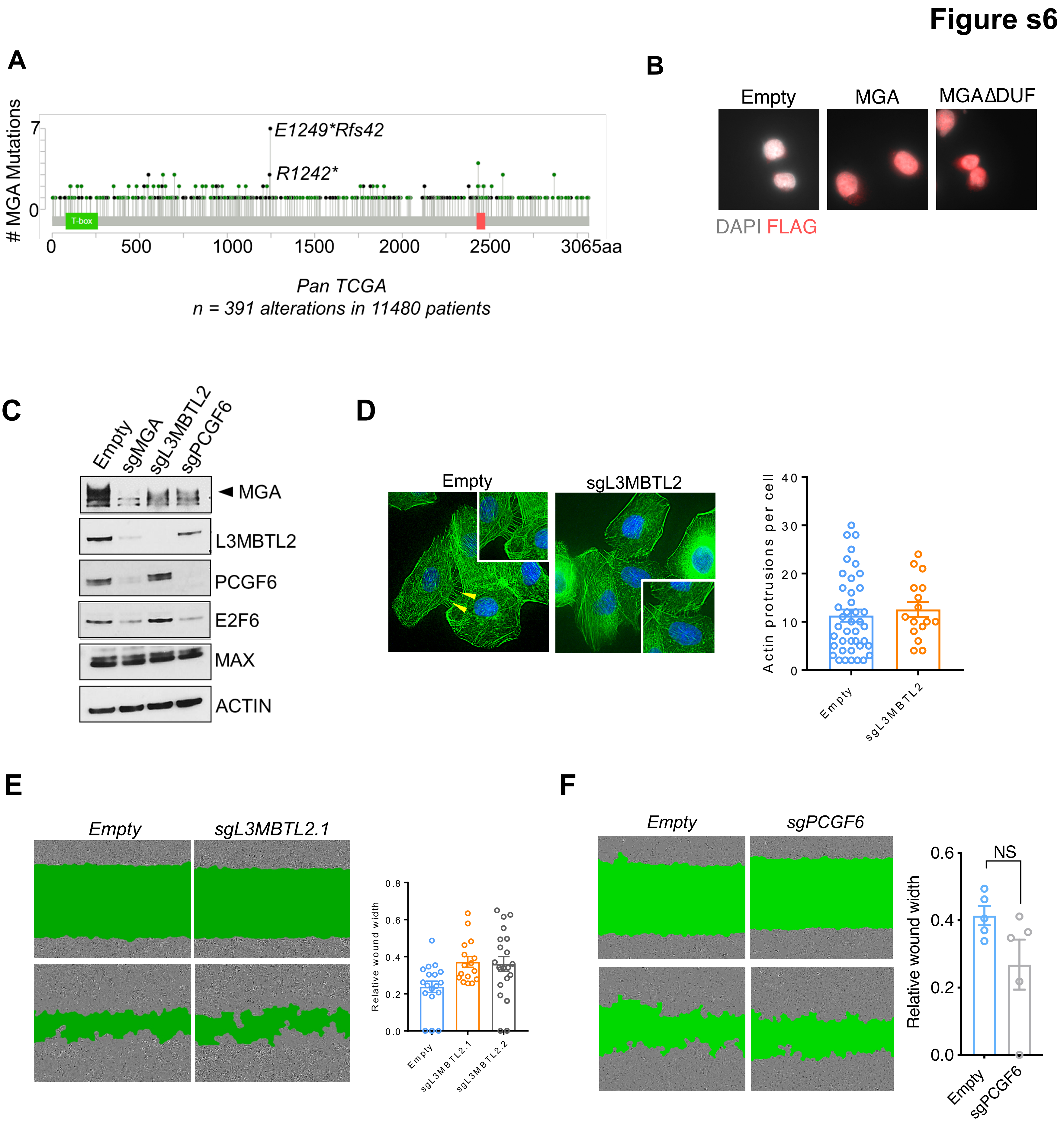


**Figure S6 The region containing the DUF4801 domain is critical for MGA's tumor suppressive function** (A) Frequently occurring mutations in the DUF domain containing region in MGA across pan-TCGA. (B) Immunostaining for FLAG in FLAG-MGA and FLAG-DUF mutant expressing cells. (C) Western blot for MGA, L3MBTL2, E2F6 and PCGF6 in sgMGA, sgL3MBTL2, sgPCGF6 and control A549 cells. (D) Actin protrusions and (E) scratch wound assay comparing sgL3MBTL2 and Empty A549 cells. (F) Scratch wound assays comparing sgPCGF6 and Empty A549 cells. Error bars represent SEM.

**Figure S7**

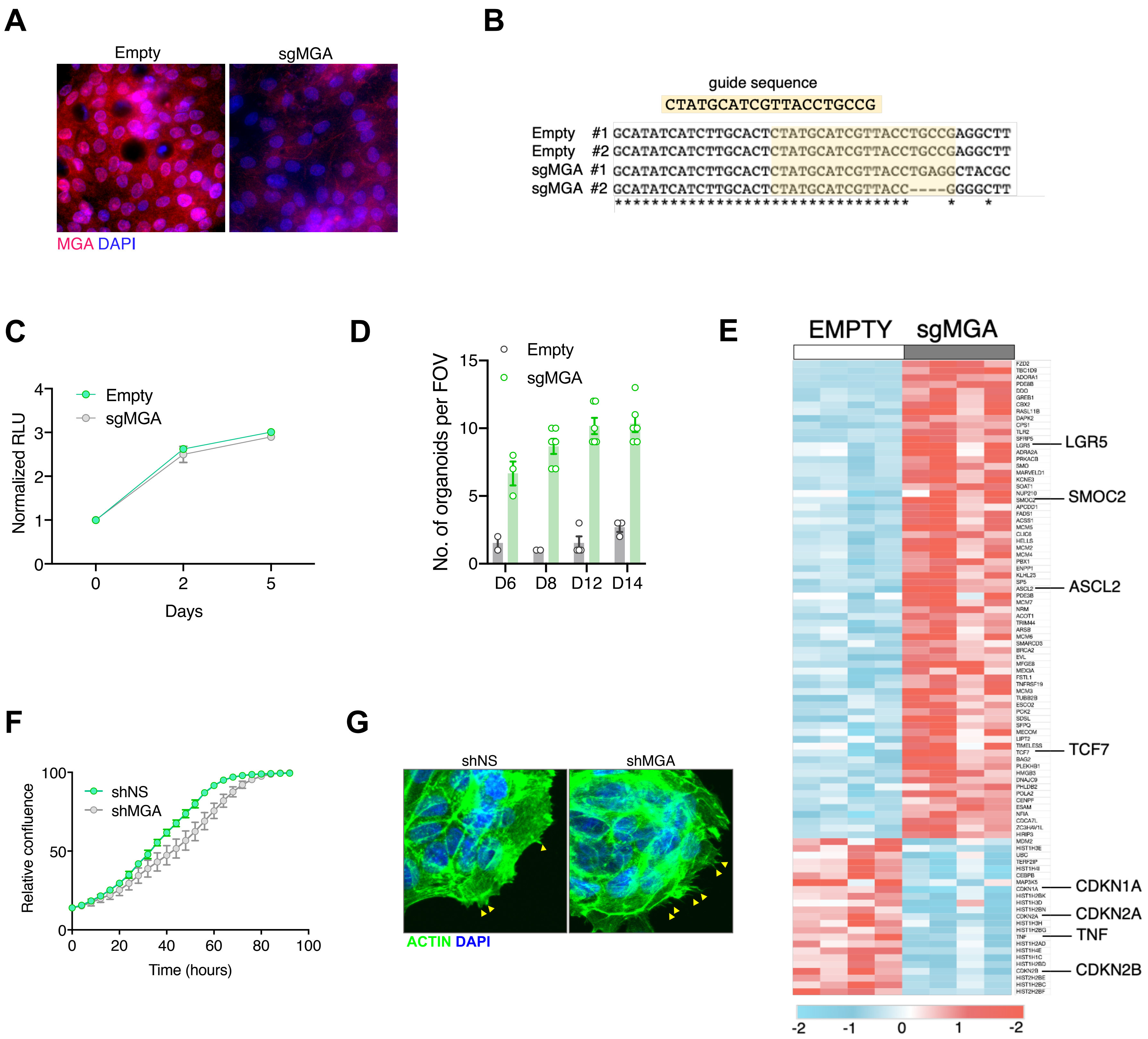


**Figure S7: MGA has tumor suppressive functions in colon cancer** (A) Immunofluorescent staining for MGA in 2D organoid cultures. (B) Sequencing to show indel formation in clonal sgMGA (#1 and #2) organoids compared to Empty vector transduced controls. (C) Growth of sgMGA and Empty organoids in 2D culture. (D) Timewise comparison of organoid growth of Empty and sgMGA transduced cells post single cell seeding. (E) Heatmap showing colon cancer stem cell signature genes and senescence associated genes that are differentially expressed between Empty and sgMGA colon organoids (LFC > 1.5, FDR <0.05). (F) Relative growth of shMGA and shNS DLD1 cells in 2D culture. (G) Representative images of actin protrusions in shNS and shMGA DLD1 cells. Error bars represent SEM.

**KEY RESOURCE TABLE**

| Reagent type (species) or resource | Designation | Source or reference | Identifiers | Additional information |
|---|---|---|---|---|
| Strain, strain background (M. musculus, male, female) | Kras | The Jackson Laboratory | RRID:IMSR_JAX:008179 | *Kras*$^{LSL-G12D/WT}$ |
| Strain, strain background (M. musculus, male, female) | KP (Kras p53) | The Jackson Laboratory | RRID:IMSR_JAX:008462 | *Kras*$^{LSL-G12D/WT}$ *Trp53*$^{Flox/Flox}$ K and P interbred at Fred Hutch |
| cell line (human) | A549 | ATCC | CCL-185 RRID:CVCL_0023 | Lung epithelial |
| cell line (human) | NCI-H23 | ATCC | CRL-5800 RRID:CVCL_1547 | Lung adeno-carcinoma, epithelial |
| cell line (human) | NCI-H2291 | ATCC | CRL-5939 RRID:CVCL_1546 | Lung adeno-carcinoma, epithelial |
| cell line (human) | LOU-NH-91 | Kind gift from Montse Sanchez-Cespedes | RRID:CVCL_2104 | Lung squamous cell carcinoma |
| cell line (human) | 91T | Kind gift From McGarry Houghton | Reference: Stabile et al. 2002 | Non-small cell lung cancer |
| cell line (human) | NCI-H1975 | ATCC | CRL-5908 RRID: CVCL_1511 | Lung adeno-carcinoma, epithelial |
| cell line (human) | DLD-1 | ATCC | CCL-221 RRID: CVCL_0248 | Colorectal adeno-carcinoma, epithelial |

| | | | | |
|---|---|---|---|---|
| Recombinant DNA reagent (human) | pcDNA3-FLAG-His-MGA | Yuzuru Shiio | | Plasmid to express full length MGA isoform2 |
| Recombinant DNA reagent (human) | pCDH-FLAG-His-MGA | This paper | | Lentiviral construct to express full length MGA |
| Recombinant DNA reagent (human) | pCDH-FLAG-His-MGAΔDUF | This paper | | Lentiviral construct to express MGA lacking the aa1003-1304 |
| Recombinant DNA reagent (*M. musculus*) | LentiCRISPRv2Cre | Addgene (Gift from Feldser lab) | | |
| Recombinant DNA reagent (*M.musculus*) | LentiCRISPRv2Cre sgMga1 | This paper | Insert (sgMga1 gRNA sequence): TAAGTGGAATGGTCGTTGGT | Lentiviral construct to transfect and express sgRNA |
| Recombinant DNA reagent (*M.musculus*) | LentiCRISPRv2Cre sgMga3 | This paper | Insert (sgMga3 gRNA sequence): TCAGAATTTTCAATATACGC | Lentiviral vector to express sgRNA |
| Recombinant DNA reagent (human) | lentiCRISPRv2 PURO | Addgene (Gift from Feng Zhang) | | Lentiviral vector to express sgRNA |
| Recombinant DNA reagent (human) | lentiCRISPRv2 sgMGA | This paper | Insert (sgMGA gRNA sequence): CTATGCATCGTTACCTGC CG | Lentiviral vector to express sgRNA |
| Recombinant DNA reagent (human) | lentiCRISPRv2 sgPCGF6 | This paper | Insert (sgPCGF6 gRNA sequence): GGTATGAAGACATTCTGTGA | Lentiviral vector to express sgRNA |
| Recombinant DNA reagent (human) | lentiCRISPRv2 sgL3MBTL2.1 | This paper | Insert (sgL3MBTL2.1 gRNA sequence): CCGGAGTTATAACAGCAGTG | Lentiviral vector to express |

| | | | | |
|---|---|---|---|---|
| | | | | sgRNA |
| Recombinant DNA reagent (human) | lentiCRISPRv2 sgL3MBTL2.1 | This paper | Insert (sgL3MBTL2.2 gRNA sequence): TTCACTTGACTCCTCCAGAT | Lentiviral vector to express sgRNA |
| Recombinant DNA reagent (human) | pLKO non-silencing | SIGMA Mission shRNA | | Lentiviral vector to express shRNA |
| Recombinant DNA reagent (human) | pLKO shMGA | SIGMA Mission shRNA | TRCN0000237941 (ID RNAi consortium) | Lentiviral vector to express shRNA |
| antibody | MGA (rabbit, polyclonal) | Novus Biological | Catalog no. NBP1-94031 | WB (1:500 in 3%BSA), IF (acetone fixation and 1:100) |
| antibody | MGA (mouse, mococlonal) | Developed at Fred Hutch | | WB (1:500 in 5% milk) |
| antibody | MGA (rabbit, polyclonal) | Kind gift from Suske Lab | | ChIP, CUT&RUN |
| antibody | Ki67 (rabbit, polyclonal) | Abcam | Catalog no. ab16667 | IF and IHC (1:200) |
| antibody | MECA32 (mouse, monoclonal) | DSHB | Catalog no. MECA-32-S | IHC (1:100) |
| antibody | MYC (rabbit, polyclonal) | Cell Signaling | Catalog no. 13987S | ChIP, CUT&RUN |
| antibody | MAX (rabbit, polyclonal) | Proteintech | Catalog no. 10426-1-AP | WB (1:1000), ChIP, CUT&RUN |
| antibody | L3MBTL2 (rabbit, polyclonal) | Active Motif | Catalog no. 39569 | WB (1:1000), ChIP, CUT&RUN |

| | | | | |
|---|---|---|---|---|
| antibody | E2F6<br>(rabbit, polyclonal) | Abcam | Catalog no. ab53061 | WB (1:1000), ChIP |
| antibody | PCGF6<br>(rabbit, polyclonal) | Proteintech | Catalog no. 24103-1-AP | WB (1:1000) |
| antibody | Rabbit IgG | Cell Signaling | Catalog. No: 2729 | ChIP |
| antibody | GFP<br>(rabbit, monoclonal) | Cell Signaling | Catalog. No: 2956S | CUT&RUN |
| sequence-based reagent (human) | siSTAG3 | Qiagen | siRNA<br>(Catalog no. SI00734440) | |
| sequence-based reagent (human) | siPODXL2 | Qiagen | siRNA<br>(Catalog no. SI04142495) | |
| software | Biorender | | Biorender.com | Used for illustrations |

Table s1: Real time primers (mouse) used in the study

| Gene | Forward | Reverse |
|---|---|---|
| Rpl4 (housekeeping) | CCGTCCCCTCATATCGGTGTA | GCATAGGGCTGTCTGTTGTTTTT |
| Podxl2 | CCCACTCAAGATGTCCTTTCC | TCACCAGCACTACAAAGAGC |
| Gpc2 | TCTTACTCGGCTTACTTCAACTG | GCTTCGCTGACCACATTTC |
| Stag3 | TCCAGTTGAGTCTGCACAAAG | CCCCTCTATGTCCTCTTGATTC |
| Snai1 | GTGAAGAGATACCAGTGCCAG | AAG ATG CCA GCG AGG ATG |
| Itga1 | GACAGCCCTTGGAATAGACAC | CTC ACA GTC TTG GAT GAC CTG |
| Snai2 | ACACATTAGAACTCACACTGGG | TGG AGA AGG TTT TGG AGC AG |
| Tdrd1 | TTCTGCTCTGTCAAGGTCATG | TTC ACT CCA TAC CCC ATT TCC |

Table S2: Real time primers (human) used in the study

| Gene | Forward | Reverse |
|---|---|---|
| GUSB (housekeeping) | CCTGCGTGTCCCTTCCTC | CGTTCTGGTCTGCCGTGAA |
| PODXL2 | CCCAGCGAAGAGAATGAAGAG | AATGGAACCTGCCTTCTCAG |
| STAG3 | GAGTGGTACATCGTCATAGCC | AATCCTGCATCCTGGTCTTG |
